# Dynamic Vision Training Transfers Positively to Batting Performance Among Collegiate Baseball Batters

**DOI:** 10.1101/2020.02.12.945824

**Authors:** Sicong Liu, Lyndsey M. Ferris, Susan Hilbig, Edem Asamoa, John L. LaRue, Don Lyon, Katie Connolly, Nicholas Port, L. Gregory Appelbaum

## Abstract

A growing body of evidence demonstrates visual, perceptual, and oculomotor abilities contribute to batting performance in baseball and there is interest in whether training such abilities can transfer positively to batting performance. The current study tested this question through a pre-registered randomized, and placebo-controlled intervention, conducted with 24 collegiate baseball players at two Division 1 universities. Athletes were randomized to receive either dynamic vision training consisting of stroboscopic, anticipatory timing, and eye quickness drills, or placebo drills stylized after control procedures in previous vision therapy studies. Generalized near-transfer was tested via a digital visual-motor task battery (*n* = 20), while sports-specific intermediate and far transfer of training were evaluated through instrumented batting practice metrics (*n* = 14) and box score performance in NCAA-sanctioned games (*n* = 12). The effects of training group were tested on these outcome measures while controlling for covariates such as pre-training expectations and site. Participants averaged 8.50 hours of training with no significant training group differences in adherence, expectations, or baseline assessments. ANCOVA revealed no group differences in measures of visual-motor skills or NCAA game statistics. However, batting practice demonstrated significant improvements in launch angle (*p* = 0.002, Cohen’s *d* = 0.74) and hit distance (*p* < 0.001, Cohen’s *d* = 0.70) for the active cohort relative to the placebo control. This controlled and pre-registered pilot study therefore provides preliminary evidence that vision training may improve batting practice performance, creating new opportunities for the transfer of skill training and warranting further study.

**Pre-Registration:** At Open Science Framework on August 9, 2018. DOI 10.17605/OSF.IO/496RX. Accessible at https://osf.io/jxh8u

## 1. Introduction

The act of hitting a pitched baseball is widely considered to be among the most challenging activities in all of sports. In milliseconds, batters must see and decipher a pitch, project its trajectory, decide whether to swing, and coordinate the movement and timing of a 2.25-inch diameter bat to contact a 3-inch ball that can be moving up to 100 mph. With such extreme demands placed on vision, it is natural to conclude that visual abilities may serve as a limiting factor in batting performance. In fact, recent literature has shown a growing body of evidence that demonstrates visual, perceptual, and oculomotor abilities are enhanced in baseball batters [1–4] and contribute to batting performance [5–9]. A strong interest thus lies in developing training programs that can produce positive transfer of visual-skill learning to on-field batting performance.

In the current pre-registered (https://osf.io/jxh8u), randomized, placebo-controlled study, we test such a vision-based training program, involving 24 collegiate baseball batters at two National Collegiate Athletic Association (NCAA) Division 1 universities. To do so, we follow a comprehensive conceptual framework of skill training and transfer [10], leverage new training opportunities with digital tools [11], and focus on testing the positive transfer to baseball batting performance.

### 1.1. Modified Perceptual Training Framework

The *modified perceptual training framework* (MPTF) [10] represents a recent theoretical framework to guide vision-based sports training approaches. These approaches typically feature vision and perceptual-cognitive training activities to enhance perceptual skills transferrable to optimal sport performance. Following principles from rep*resentative learning designs* (RLD) [12], MPTF helps classify a given training approach along three continuous dimensions, including the similarity between training and sports-defined (1) skills, (2) stimuli, and (3) responses. MPTF further predicts that training programs maximizing correspondence along these dimensions will result in the greatest transfer to improvement on the playing field. As such, MPTF offers a promising guide for designing or refining existing sports training programs, particularly those with a core interest of transferring training benefits to real-world sports performance.

Two categories of sport-specific training following the MPTF are perceptual-cognitive skill training and sports vision training. *Perceptual-Cognitive Training* aims to sharpen skills, such as anticipation and decision-making, with sports-specific video footage or images in tasks that adopt temporal and/or spatial occlusion paradigms [13]. In contrast, *Sports Vision Training* (also referred to as generalized vision training) focuses on strengthening oculomotor and visual-perceptual skills through generic visual stimuli (e.g., shapes, visual fields, depth cues) in customized vision-therapy-based tasks [14]. As it focuses primarily on oculomotor and visual-perceptual skills, sports vision training generally locates lower than perceptual-cognitive training on all the three MPTF dimensions, leading to less evidence in support of transfer to sports performance [15]. Despite this, there are studies demonstrating evidence that sport vision therapy can produce improvements in sport performance [16].

### 1.2. Skill Training and Transfer

*Skill transfer* refers to either a gain (i.e., positive transfer) or loss (i.e., negative transfer) in performance on a criterion task resulting from practicing a training task [17]. The transfer of learned skills is a century-old enterprise (reviewed in [18, 19]) garnering renewed interest in light of the recently increased prevalence of ‘brain training games’ [20]. Learned skills transfer is based largely on the *theory of identical elements* (TIE). TIE contends the degree to which training transfers depends on the proportion of shared skill elements between training and performance tasks [21]. Within the transfer literature there is substantial evidence in support of the *near transfer* of skill training, wherein practicing one task benefits performance on another task that shares a high proportion of skill elements [17, 22]. *Far transfer*, in which learning projects from training situations that share fewer skill elements with the desired performance task, is less prevalent, particularly as it relates to sports [23, 24].

According to contemporary taxonomy, transfer depends on both the *content* of what is transferred and the *context* in which training and transfer happen [22]. Content addresses the specificity-generality of transfer, whereas context helps determine how far the transfer carries through criteria involving task modality (e.g., visual versus auditory) and functional domain (e.g., physical, temporal, social) contexts. Viewing these dimensions from the sports training perspective, the content-context transfer taxonomy (CCTT) agrees fundamentally with MPTF and RLD. For transfer to occur, the sports training program should sufficiently simulate the sports performance environment by presenting corresponding stimuli, demanding corresponding responses, and improving the most relevant skill elements.

### 1.3. New Opportunities for Baseball

Following the MPTF and CCTT, recent innovations have opened new opportunities for developing vision-based training to improve sports performance in general, and baseball performance in particular. A fast growth in new technologies utilizing digital tools now provides several novel approaches for training visual skills [11]. Many of these approaches can be deployed during natural training activities to promote greater skill, stimulus, and response correspondence, as highlighted in MPTF. For example, the use of stroboscopic eyewear creates situations where individuals can train under intermittent visual conditions and has now been tested in a number of controlled [25, 26] and pseudo-controlled studies [27–30], providing evidence that this type of training can improve aspects of vision and motor control that may transfer to enhanced on-field performance (reviewed in [31]).

These technologies have also increased accessibility to robust and meaningful sports data, and helped identify visual-motor skills associated with improved athletic performance. For instance, with a range of vision screening, eye tracking, and pitch-by-pitch baseball batting data in professional batters, it was recently shown better eye-tracking-based oculomotor skills predict improved plate discipline performance (e.g. lower propensities to swing at pitches outside of the strike zone) and increased the highest league level attained [32]. Similarly, another recent study linked superior visual-motor skills to higher on-base percentage, and lower walk and strikeout rates in professional baseball players [5]. Other reports have demonstrated superior baseball statistical production is correlated with better eye-hand reaction times [6], more precise dynamic stereoacuity [33], and quicker visual reaction and recognition times [20, 34], thereby adding to the evidence that visual skills contribute to batting performance.

Beyond correlational findings, past studies have also contributed to the understanding of vision-based training in interceptive sports, including cricket [35], softball [36, 37], field hockey [38], tennis [39], table tennis [40], and baseball [16]. Despite some disagreements [41], this body of work has generally supported the utility of vision-based training for improving visual skills in interceptive sports. Some vision-based training protocols [16,42] were even argued to generate positive far transfer to sports-specific performance. However, such causal inferences must be interpreted with caution given the presence of several inferential threats, such as lack of matched placebo control groups, the absence of randomization, and no limitation on researcher degree-of-freedom through pre-registered designs and analyses [43]. As such, clinical-trial-like training study addressing these limitations is needed to further advance the field and determine if visual skill training can transfer to improved sport performance [44].

### 1.4. The Present Study

Guided by the MPTF and broader skill transfer literature described above, this study took advantage of new training opportunities made available via digital tools and aimed to explore innovative vision-based protocols with the capacity to transfer to baseball batting performance. As such, this work focused on testing meaningful transfer benefits to real-world sports performance, while striving for the high methodological standards of a pre-registered [45], randomized, and placebo-controlled study. Moreover, this study attempted to maximize the sample size of expert participants by conducting the protocols with baseball batters from two NCAA Division 1 collegiate teams in parallel.

Skills likely to contribute to baseball batting performance were identified from past correlational and training evidence, including dynamic vision (object-tracking, accommodation, and vergence), oculomotor speed/accuracy, anticipatory timing, and visual attention [5, 32, 46, 47]. These represent the targeted skill elements, and collectively possess a high correspondence between training and sports-defined skillset according to MPTF. Stations were established to train the targeted skills using three digital tools, including a tablet-phone pair, an 18-foot long LED light rail, and stroboscopic eyewear. The latter two training stations were designed to maximize the stimulus and response correspondence outlined in MPTF. The light rail was predominantly used to simulate a baseball pitch and was set up in an according spatial manner. Stroboscopic glasses were used in a set of drills, taking the form of simple and sports-atypical practice at the start (to foster tool familiarization), then transitioning to more sports-specific drills like catching and controlled hitting in later training phases. Placebo-control training incorporated similar oculomotor, reaction, and visual-perceptual tasks modeled after traditional vision therapy protocols, but altered such that they did not actually engage those skills.

To test the effects of this controlled intervention, pre- and post-training measures of participants’ instrumented batting practice performance and NCAA-sanctioned game statistics were obtained, along with six non-targeted visual-motor skills. Based on CCTT, the six visual-motor skills were considered measures of *generalized near transfer* because they tested the transfer-of-learning to visual-motor abilities measured with similar stimuli and on digital equipment identical to the training activities, but did not replicate the training tasks. Instrumented batting practice represented *sports-specific intermediate transfer* because it focused on generalization to baseball batting, but was not directly targeted in the study’s training activities. Season game statistics were treated as *sports-specific far transfer* measures due to the complete generalization to competitive contexts. As described in the pre-registration for this study [45], it was hypothesized that the active vision training protocol would lead to positive transfer in each domain, relative to the placebo control, resulting in improved visual-motor skills and better metrics of batting practice and game production.

## 2. Methods

### 2.1. Participants

Participants consisted of NCAA Division 1 baseball players from Duke University (Duke) and Indiana University, Bloomington (IU). Initially, 26 individuals expressed interest and were assessed for eligibility, with two individuals being excluded due to unrelated injuries. As a result, a total of 24 participants (14 from Duke and 10 from IU) were enrolled and provided informed consent following the standards approved by each university’s research ethics committees that each conformed to the tenants of the Declaration of Helsinki.

### 2.2. General Study Procedures and Timeline

This study was reviewed and approved by the Institutional Review Board of each respective institution. This involved gaining approval for data sharing of anonymized databases between institutions. As such, each participant was assigned a generic alphanumeric identifier to be associated with all study-obtained data to maintain confidentiality. Study-related activities began with informed consent, proceeded until the post-training assessments, and concluded with payment at the rate of $40/hour of participation at IU and $20/hour at Duke.

All study procedures followed a pre-registration that was reviewed and published before the start of study activities. The study timeline, activities, and participant flow are outlined in Figure 1 and described in detail in the sections below. Pre-training evaluations were initiated in September 2018 and post-training evaluations were completed by January 2019. Batting practice occurred several times a week over the months of September through November at Duke and took place in two 10-day intervals at the beginning and end of this time period at IU. Regular season game statistics from the 2018 and 2019 collegiate baseball season (excluding exhibition games and playoffs) were acquired over the months of February through June for all participants who played sanctioned regular-season games during those seasons.

**Figure 1.**
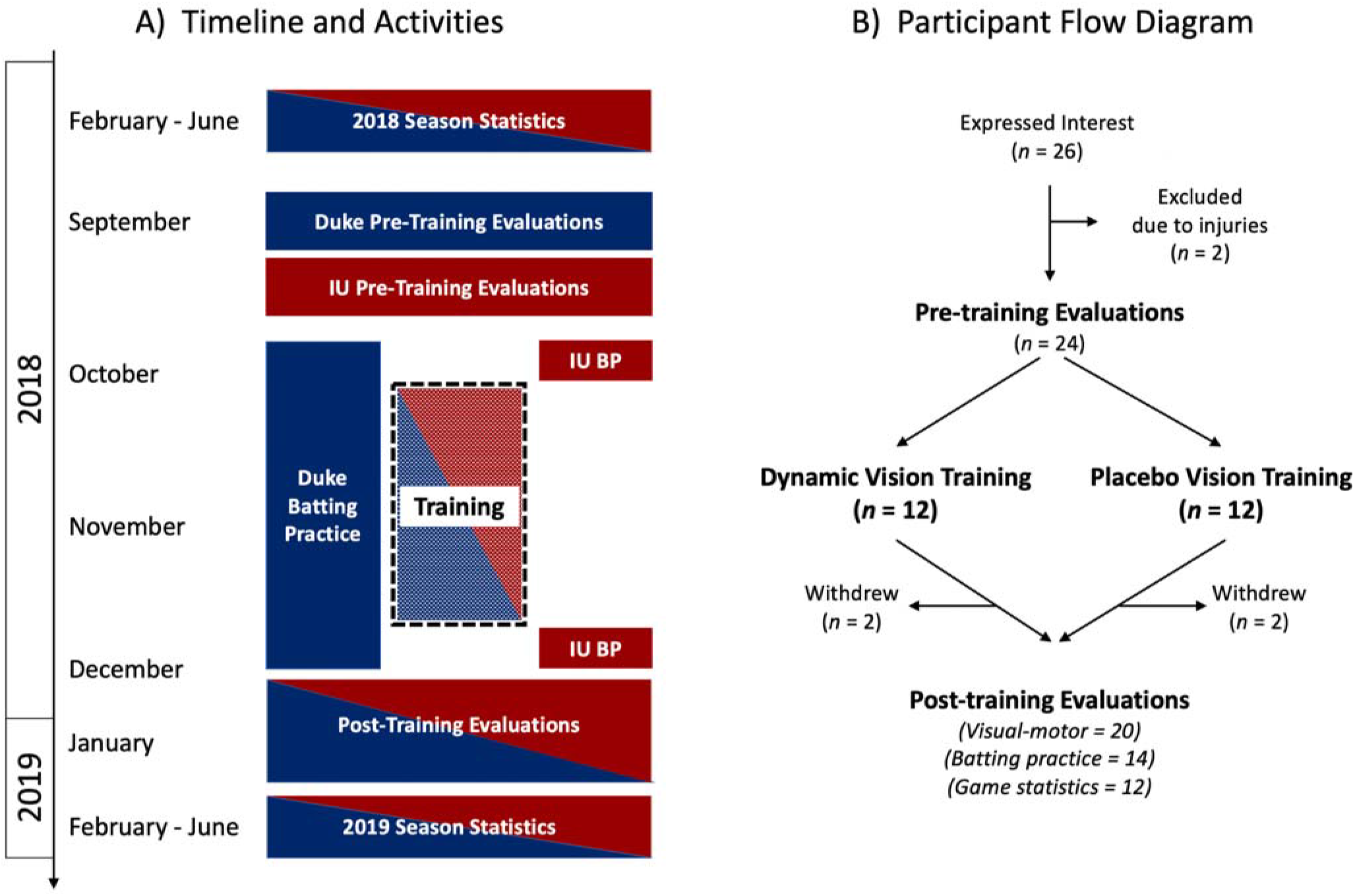
A) Illustration of study timeline and activities. Activities conducted at Duke are colored in dark blue, while activities conducted at IU are colored in red. B) Illustration of participant flow through the study activities with counts and attrition. The y-axis to the left represents time and applies to both panel A and B.

All participants enrolled at the outset of the study received pseudo-random assignment into one of the two training arms: either the active vision training group referred to as <u>*Dynamic Vision Training*</u> (DVT) or a placebo control training group, <u>*Placebo Vision Training*</u> (PVT). Group assignment was done differently at the two sites. At Duke, time slots were filled first according to the athletes’ availability, then group assignments were made randomly among all participants to maximize equipment use. At IU, enrolled batters were first split in two groups: (A) returning players who regularly batted in the previous year, and (B) new players or players who did not regularly bat in the previous season. Each batter within group A and B was then randomly assigned to either the DVT or PVT groups. This avoided the possibility of randomly including all returning players into one group or the other. All participants underwent training activities in 30-minute sessions (described in Section 2.3) across and up to 10 weeks within the assessment period. Four participants (two from Duke and two from IU) withdrew during the training program due to schedule conflicts. As such, 20 participants completed the full training protocol and the post-training visual-motor assessments.

Three types of outcome measures were collected: *visual-motor evaluations, instrumented batting practice*, and *NCAA season game statistics* (see Section 2.4). The evaluation of visual-motor abilities using the Senaptec Sensory Station Tablet represented *‘generalized near-transfer’* outcome measures and were collected on all 20 participants who completed the training (12 from Duke and eight from IU). Instrumented batting practice represented *‘sports-specific intermediate-transfer’* outcomes and were available for 14 participants (six from Duke and eight from IU). NCAA game statistics represented *‘sports-specific far-transfer’* outcomes, which were available for 12 participants (six from Duke and six from IU).

### 2.3. Training Procedures

Each training session lasted approximately 30 minutes with individuals proceeding through a designated set of activities defined according to procedures for each training group. These training activities were conducted in an adaptive format, such that the difficulty of each drill was customized based on an individual’s previous performance according to pre-established advancement criteria (detailed in the **Supplement 1**). While training sessions took place from late September 2018 through early December 2018, no single participant took part in more than 10 weeks of training. The total number of visits was dictated by participants’ availability and ranged from 7 to 22 sessions at Duke and 17 to 25 at IU, leading to an overall mean of 17 visits among the participants who completed the study.

At Duke, training took place in indoor facilities located next to the baseball training field and in the team’s covered batting cages. At IU, training took place in research facilities located at the Lawrence D. Rink Center for Sports Medicine and Technology. Authors SL, SH, EA, JL, LGA served as trainers across the sessions at Duke, while authors LMF, NLP, DL, KC, PK, and collaborating students from the IU School of Optometry served as trainers at IU.

#### 2.3.1. Active Cohort – Dynamic Vision Training (DVT)

DVT consisted of three types of drills: (1) stroboscopic vision training using the Senaptec (Senaptec LLC, Beaverton, OR) Stroboscopic eyewear, (2) oculomotor and anticipatory timing training using the Senaptec Synchrony light rail, and (3) dynamic vision training using the Senaptec Sensory Station Tablet. Approximately 10 minutes of each training session were dedicated to each type of drill, with activities arranged in brief circuit training routines. General descriptions of the training activities are listed below and illustrated in Figure 2A, while detailed descriptions and procedural criteria can be found in **Supplement 1**.

**Figure 2.**
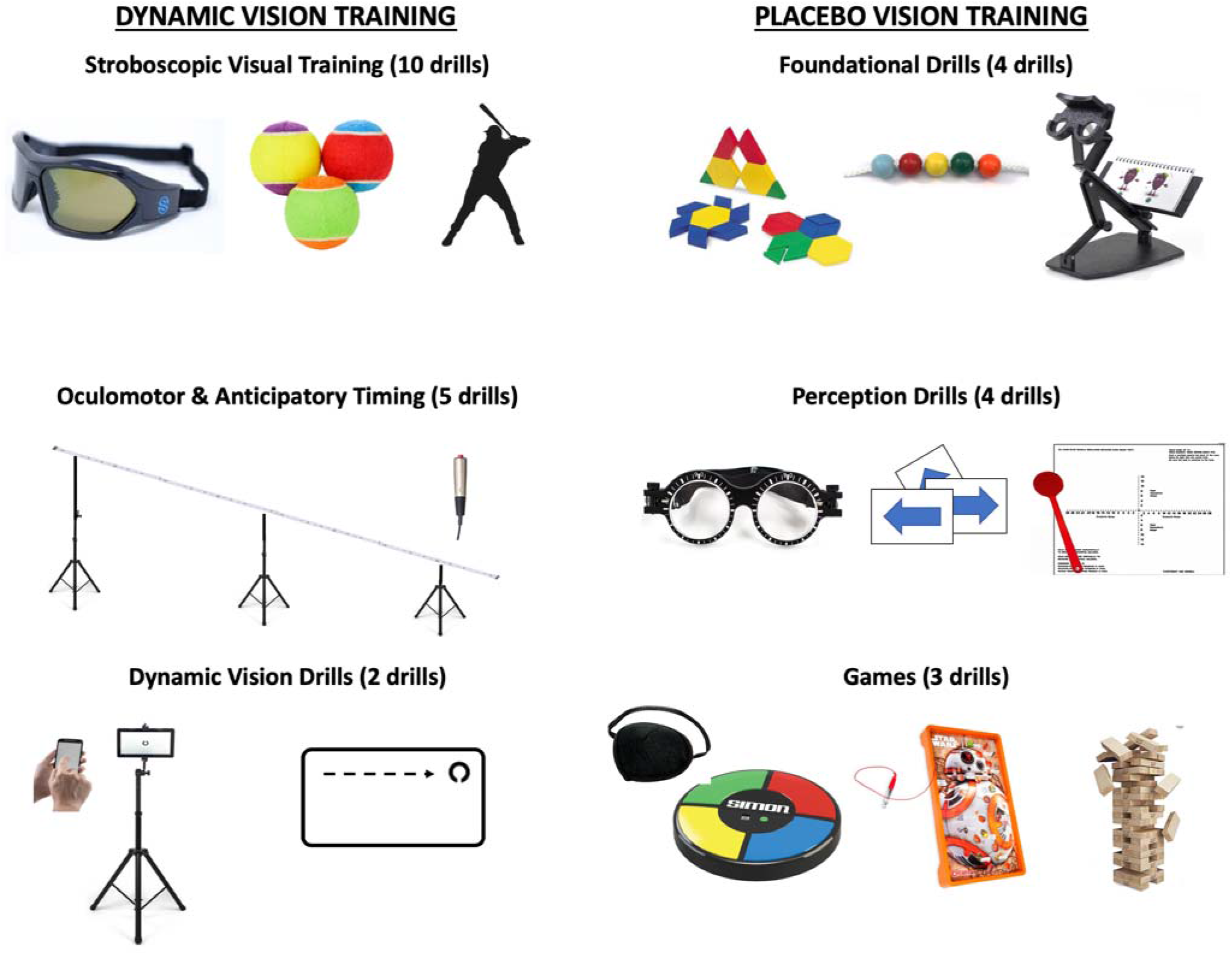
Illustrations and examples of the Dynamic Vision Training or Placebo Vision Training activities. DVT consisted of 10 different Stroboscopic Visual Training drills (which involved wearing strobe eyewear during catching and hitting drills), Oculomotor and Anticipatory Timing drills (which required participants to track or respond to fast moving lights on a light rail), and Dynamic Vision drills (completed on a digital tablet). PVT consisted of Foundational Drills (including vertical oculomotor stability, pattern recognition, and attention to detail), Perception drills (with adapted visual therapies) and Games (which incorporated games such as Simon™, Operation™, and Jenga™ with visual challenges such as eye patches or diopter lenses). PVT drills were modified such as not to directly engage or train horizontal oculomotor skills, visual-reaction timing, or depth/field of view perception.

- **Stroboscopic Visual Training** included 10 specific drills, ordered from simple and sport-general (such as tossing and catching a ball) to difficult and sport-specific (such as batting a pitched ball). Participants wore strobe eyewear which allowed for temporal phasing between opaque and transparent lenses at levels of variable ratio ranging from 1 to 8 as controlled by a button on the right temple of the frame. At all levels, the transparent state lasted 100 ms in each cycle, with longer periods of occlusion at higher levels. These ranged from 67 ms at level 1 to 900 ms at level 8. Each participant escalated through the levels at an individual pace according to criteria defined by performance on each task, for example, leveling up after successfully catching the ball nine times out of ten.
- **Oculomotor and Anticipatory Timing** Training included five specific drills using the 18-foot long Synchrony light rail. This rail includes 100 discrete LED lights with a handheld response button and was positioned to mimic the trajectory of a baseball pitch (ramping downward from 6.50-feet tall to 3-feet tall over an 18-foot span). The first two drills focused on oculomotor skills of smooth pursuit and saccadic movement, with the participant standing perpendicular to the rail and moving only his eyes to follow the continuous movement of the light across LEDs (smooth pursuit) or jump to the location of lights as they appeared at new, random locations across the rail (saccade). The remaining three centered on visual-motor skills of anticipatory timing and prepotent response inhibition, with the participant standing at the short end of the rail and holding a response trigger. The difficulty of the drills was generally controlled by adjusting the movement speed of visual stimuli on the light rail.
- **Dynamic Vision Drills** included two training drills: Near-Far Quickness and Dynamic Visual Acuity. The Near-Far Quickness drill aimed to train the speed and accuracy of accommodative vergence eye movements by alternating targets between near (1 ft) and far (10 ft) distances. Successive strings of three stimulus units such as Landolt Cs were shown on the far and near screens. Participants swiped the handheld screen in the direction of the gaps of the two similar Landolt Cs, which is inconsistent with the third Landolt C, before the next stimuli was presented on the other screen. Scores were determined as the total number of correct targets identified in 30-seconds. Difficulty on this task was controlled by elevating the passing criteria while diminishing the visual contrast between targets and background. The Dynamic Visual Acuity drill aimed to train acuity for rapidly moving targets by presenting Landolt C stimuli that moved rapidly from one corner of the far screen (10 ft) to another. Participants were instructed to swipe the handheld controller in the direction of the gap in the C as they perceived it during the movement path of the stimulus. Difficulty was controlled by increasing movement speed and duration of visibility for the Landolt stimulus.

#### 2.3.2. Placebo Cohort – Placebo Vision Training (PVT)

The placebo control methodology was modeled after control conditions in previous vision therapy trials, particularly the Convergence Insufficiency Treatment Trial [48]. Drills were adapted from traditional vision therapy protocols but were modified so as not to engage the traditional targeted skill. For example, the Brock String procedure engages a patient’s binocular vision system to increase vergence and divergence skills without suppression. This task requires the use of both eyes to engage these systems. However, PVT participants performed this exercise with an eye-patch, negating any visual feedback and thus limiting any vergence training effects.

Each session consisted of three categories of drills: (1) Foundational Drills which aimed to consistently provide a clear metric against which participants could improve in order to maintain engagement, (2) Perception Drills which utilized vision therapy techniques in order to contribute scientific legitimacy to the control, and (3) Games paired with placebo vision therapy techniques. On average, each PVT session included a subset of eight drills, allowing participants to perceive increased difficulty as tasks changed over time. Time spent with each test varied as participants progressed through training, but each session totaled 30 minutes. General descriptions of the training activities are listed below, while detailed procedures can be found in **Supplement 1**.

- **Foundational Drills** included five specific drills from traditional vision therapy protocols redesigned to mimic engagement of the oculomotor and binocular vision systems while still appearing as legitimate, “old school” vision therapy techniques. These included stick-and-straw (fixation, vertical versions), unilateral Ductions (horizontal and vertical versions), unilateral Bernell-o-Scope (fixation, tracing), unilateral Brock’s String (fixation), and Modified Thorington (fixation, phoria). Instructions varied from using one or both eyes, to wearing plus-powered fogging lenses to give the appearance of scaling for difficulty.
- **Perception Drills** included three specific drills utilized in traditional vision therapy and designed to engage the static visual-perceptive skills of participants. These included parquetry block patterns (fixation, attention, problem-solving), visual memory (fixation, working memory), and visual illusions such as the Necker Cube, Lilac Chaser, and Motion Induced Blindness (fixation, attention)^49^.
- **Games** included three drills, each of which applied placebo vision therapy techniques to classic games such including Jenga (fixation, hand-eye control), Operation (fixation, hand-eye control), or red/green suppression glasses with normal playing cards (fixation). Difficulty was increased by using an eye patch or fogging lens to block the use of one eye and create the illusion of training.

### 2.4. Measured Variables

#### 2.4.1. Visual-Motor Evaluations – Generalized Near Transfer

To assess generalized near transfer of sensorimotor learning from the training program, participants were evaluated on six visual-motor tasks using the Senaptec Sensory Station Tablet. These tasks were derived from the Nike LLC (Beaverton, OR) Sensory Station device, which has been used extensively to develop sports specific normative data. These authors, and others, have studied variability associated with psychometric performance on these tasks [50] and shown predictive relationships with batting [5,32] and hockey performance [51].

The device was set up indoors, with the tablet screen attached to an adjustable tripod which allowed the screen to be placed at eye level for each participant. Depending on the activity, participants were asked to stand either 10 ft or 2 ft away from the tablet. For certain activities, the participant held a smart phone device approximately 40 cm away from their face and ~30° below eye level as a response remote for the task. This evaluation session took less than 30 minutes and consisted of the six following tasks:

- **Visual Clarity** is a measure of static visual acuity obtained by having participants report the orientation of gaps in a Landolt C ring, presented at distance, and adjusted in size according to an adaptive staircase procedure. Scores are reported in LogMAR units with smaller values indicating better performance.
- **Contrast Sensitivity** measures the minimal foreground contrast shown in static targets displayed at distance. Stimuli are presented at 18 cycles-per-degree and adjusted in contrast according to an adaptive staircase procedure. Results are reported in log contrast with larger values indicating better performance.
- **Near-Far Quickness** is a measure of how quickly participants can visually accommodate back and forth between near and far visual targets in 30 seconds, without sacrificing response accuracy. Scores indicate the number of correctly reported targets with larger values indicating better performance.
- **Multiple Object Tracking** is a measure of how well participants can maintain accurate spatial tracking of multiple moving targets according to an adaptive staircase schedule. Scores are computed as a composite of movement speed thresholds and tracking capacity, with larger values indicating better performance.
- **Perception Span** is a measure of spatial working memory derived by having participants recreate the locations of briefly presented targets that are flashed in a grid of circles. The number of targets and the size of the grid increases with correct responses, and the final score indicates the combined total of correct responses, minus errors, across all levels. Larger values indicate better performance.
- **Reaction Time** is the elapsed time between when rings on the touch pad change color, and when participants can remove their index finger from the touch sensitive screen. Scores are reported in seconds with smaller values indicating better performance.

Duke participants were each tested once on the pre- and post-training evaluations, while IU participants were assessed five times at each occasion of pre- and post- training evaluations.

#### 2.4.2. Instrumented Batting Practice – Sports-Specific Far Transfer

To obtain controlled measures of batting performance, quantitative pitch-by-pitch tracking was recorded during instrumented batting practice conducted at each study site. Due to logistic and equipment constraints, the procedures of batting practice and the quantity of the data differed at the two sites. As illustrated in Figure 3A, each of the eight participants at IU completed about 200 trials of batting practice facing pitches projected by a pitching machine (*M* = 75.03 mph, *SD* = 2.63 mph) at a distance of approximately 60 ft both prior to and after the training period (with a total of 3237 observed pitches, prior to exclusion). The batting practice performance at Indiana University was tracked and recorded by the HitTrax system (InMotion Systems LLC., Northborough, MA).

**Figure 3.**
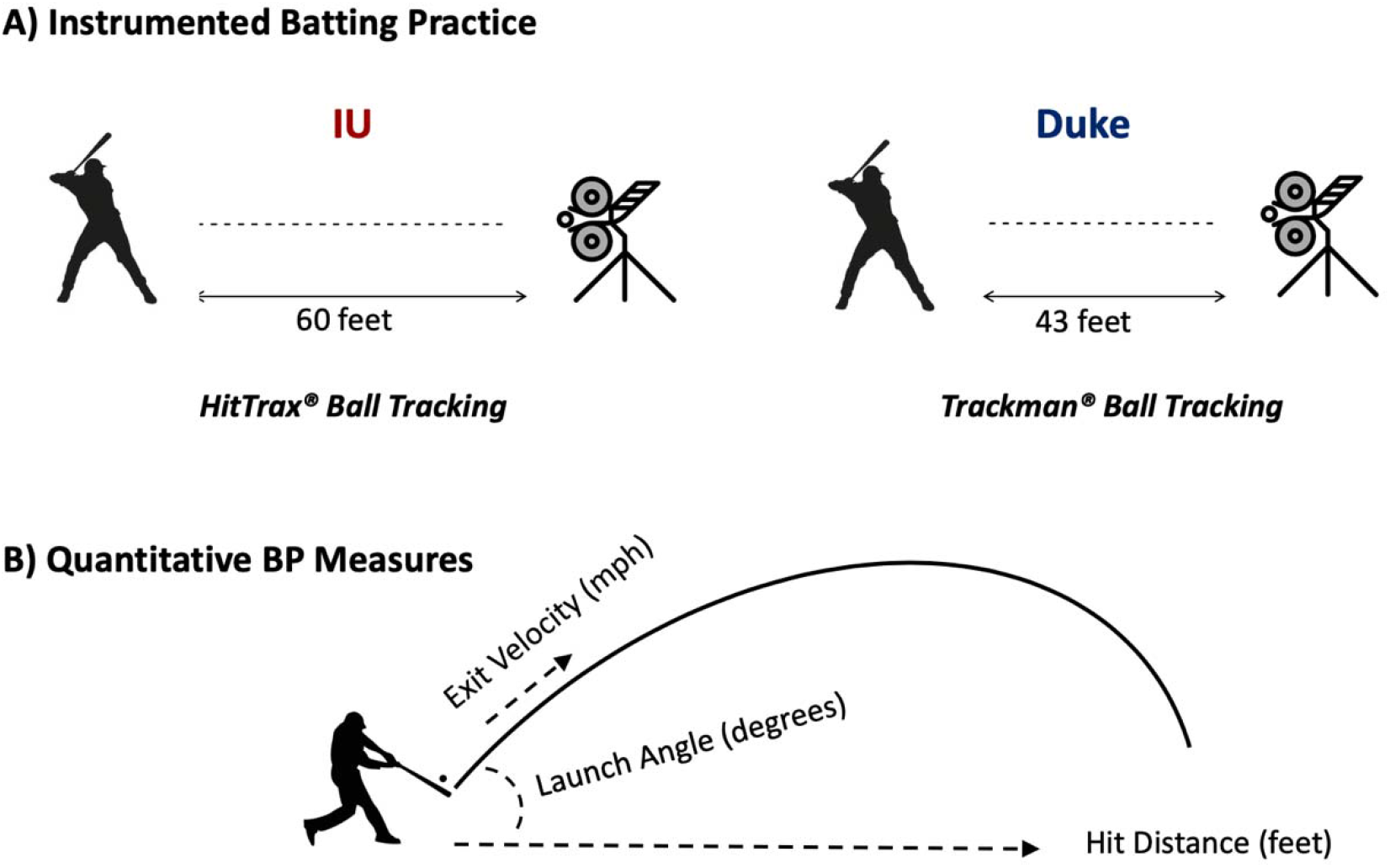
A) Illustration of instrumented batting practice at the two study sites. B) Illustration of three quantitative measures of batting practice (BP) performance; Exit Velocity in miles/hour, Launch Angle in degrees relative to the ground plane, and Hit Distance from batter to landing location of hit ball.

Batting practice at Duke was performed continuously throughout fall 2018 with the first practice sessions starting ~4 weeks prior to the start of the training period. This batting practice was performed as part of team activities with individual sessions comprised of scenario-based batting drills (for example, trying to hit specifically to advance a runner from 2^nd^ to 3^rd^ base) and ‘free-swing’ hitting drills. Only free-swing blocks are considered for analyses here. While not all Duke batters continued the batting practice throughout the study, six batters completed batting practice sessions after the end of their training activities. Therefore, pre- (*n* = 277) and post- (*n* = 145) training batting practice observations are available for these batters. These batters faced pitches (*M* = 66.74 mph, *SD* = 2.32 mph) propelled by a pitch machine at a distance of approximately 43 ft. Batting practice performance at Duke was measured and recorded by the Trackman Doppler Radar system (Trackman LLC., Stamford, CT).

As illustrated in **Figure 3B**, the following quantitative tracking performance variables were calculated for each hit at each site.

- **Exit Velocity**, representing the ball’s traveling speed (mph) immediately after bat contact.
- **Launch Angle**, representing the angular degree () of the ball’s initial flight trajectory after bat contact relative to the horizontal plane.
- **Hit Distance**, representing the distance (in feet) of the ball flight prior to touching the ground after bat contact.

To be included in subsequent analyses, individual batting practice observations needed to travel no less than 10 feet, possess a launch angle no less than −20, and possess an exit velocity no less than 50 mph. This resulted in 468 out of 3659 total possible observations (12.8%) being excluded in the final analyses.

#### 2.4.3. NCAA Game Statistics – Sports-Specific Far Transfer

The 2018 and 2019 season box scores were available for 12 participants. These records were used to generate the number of at-bat instances and to calculate a total of five season statistics:

- **Batting Average** is the rate of hits per at bat.
- **On-Base Percentage** is the rate to reach bases.
- **Slugging Percentage** is the mean number of total bases per at bat.
- **Walk Percentage** is the rate of walks per plate appearance.
- **Strikeout Percentage** is the rate of strikeouts per plate appearance.

#### 2.4.4. Training Expectation Surveys

Two Likert-type survey questions were designed to measure participants’ expectations of training benefits from the intervention based on previous research [52]. One question asked about the expected benefits of training on vision: “Do you think completing the vision training would lead to better vision in baseball?” The other question addressed expected training benefit on baseball performance: “Do you think completing the vision training would lead to better performance in baseball?” Both questions were asked with response options ranging from 0 to 6. Here, 0 indicated “no benefit”, 1 indicated “a little benefit”, and 6 indicated “a lot of benefit”. Survey questions were completed by participants both prior to and after the training intervention.

### 2.5. Analyses

Analytical procedures for this study followed the plans outlined in the pre-registration with the primary goal to test transfer of training to batting performance as represented in instrumented batting practice and NCAA season statistics. All analyses occurred at the observation level of individual trainees. When lower level observations were available for individual trainees (e.g., trial-by-trial performance in batting practice), individual means were obtained and analyzed. Four steps of analyses were performed. The first step focused on describing the data according to the outcome measures (**Figure 1**). The second step tested potential threats to the attempted causal inferences in the next two steps, including differential expectation, differential training session adherence, and baseline group differences. Two-sided *t*-tests were used in testing for differential expectation and training session adherence, and a regression model was used to test for baseline group differences. The third step answered the question of whether a multi-week training program led to generalized near transfer to improve visual-motor skills.^a^ Analysis of covariance (ANCOVA) models were fit for each of the generalized visual-motor skill measures. The fourth and final step represented the core interest of testing transfer of learning to answer the pre-registered research question: whether a multi-week training program generates positive transfer to batting performance. Again, ANCOVA models were fit to the performance variables assessed at pre- and post-training periods. For instance, pre-training Hit Distance in the instrumented batting practice served as the covariate for testing Training Group differences on post-training Hit Distance, while controlling for site and pre-training expectations. Because of differences in the equipment and procedures for the instrumented batting practice between the two sites, data from the two sites were also analyzed and reported separately.

Both complete-case and intent-to-treat analyses were performed. The intent-to-treat analysis, reported in **Supplement 2.1**, helped check the inferential threat of differential attrition in the study [53]. Missing values were imputed using the last-observation-carried-forward/backward method when at least one observation is available on a task for a given batter and were imputed using group means when no observation is available on a task for a given batter. Results from the complete-case analysis were reported while those from the intent-to-treat analysis served as part of sensitivity analysis [54]. All analyses were performed in R (R Core Group, 2019) and the *α* level was set at 0.05. Given the directional hypotheses predicting training gains in the pre-registration of the study, the *p* values associated with testing the training group differences are from one-sided tests, whereas the rest of the *p* values are from two-sided tests.

## 3. Results

### 3.1. Descriptive Information

The current study included longitudinal data consisting of training data and completed pre/post visual-motor evaluations for 20 participants, completed pre/post batting practice data for 14 participants, and NCAA game statistics for both the 2018 and 2019 season for 12 participants. **Table 1** summarizes the participant number, completed number of training sessions, the training expectation survey variables, and the six visual-motor evaluation variables (which helps testing generalized near transfer of training) according to levels of training group and site, respectively. **Table 2** summarizes the similar groups of variables with a focus on batting practice variables and NCAA season statistics (used in testing for sports-specific intermediate and far training transfer, respectively) according to levels of training group and site. Similar summaries of variables according to the factorial combinations of levels of training group and site are available in **Supplement 2.2**.

**Table 1:** Means (and SDs if applicable) for measures of participation, expectations and performance on the visual-motor evaluations

|  |  | Group |  | Team |  | All |
| --- | --- | --- | --- | --- | --- | --- |
|  |  | DVT | PVT | Duke | IU |  |
| Number of Participants |  | 10 | 10 | 12 | 8 | 20 |
| Number of Training Session |  | 17.2 (7.3) | 16.8 (6.9) | 12.7 (5.1) | 23.5 (2.8) | 17 (6.9) |
| **Expectations** |  |  |  |  |  |  |
| Vision Expectations | Pre | 5 (0.47) | 5.1 (0.88) | 5.33 (0.65) | 4.62 (0.52) | 5.05 (0.69) |
|  | Post | 5.6 (0.7) | 5.4 (0.84) | 5.67 (0.65) | 5.25 (0.89) | 5.5 (0.76) |
| Perform Expectations | Pre | 4.6 (0.84) | 5.2 (0.79) | 5.33 (0.78) | 4.25 (0.46) | 4.9 (0.85) |
|  | Post | 5 (0.67) | 5.2 (0.92) | 5.5 (0.67) | 4.5 (0.53) | 5.1 (0.79) |
| **Visual Motor Evaluations** |  |  |  |  |  |  |
| VC (LogMAR) | Pre | -0.13 (0.093) | -0.179 (0.11) | -0.133 (0.12) | -0.188 (0.043) | -0.155 (0.1) |
|  | Post | -0.172 (0.11) | -0.182 (0.078) | -0.149 (0.1) | -0.218 (0.07) | -0.177 (0.096) |
| CS (log) | Pre | 1.56 (0.19) | 1.67 (0.26) | 1.58 (0.22) | 1.67 (0.25) | 1.62 (0.23) |
|  | Post | 1.57 (0.24) | 1.75 (0.27) | 1.68 (0.31) | 1.63 (0.18) | 1.66 (0.26) |
| NFQ (score) | Pre | 32 (5.7) | 29.3 (7.2) | 29.2 (6.8) | 32.8 (5.8) | 30.7 (6.5) |
|  | Post | 37.2 (7.1) | 34.6 (4.3) | 34.5 (6.7) | 38.1 (3.6) | 35.9 (5.8) |
| PS (score) | Pre | 45 (11) | 50.4 (13) | 48.8 (13) | 45.9 (11) | 47.7 (12) |
|  | Post | 50.2 (11) | 51.7 (10) | 51.6 (12) | 50 (9.4) | 51 (11) |
| MOT (score) | Pre | 1776 (288) | 1852 (564) | 1925 (362) | 1647 (511) | 1814 (438) |
|  | Post | 1916 (493) | 1887 (650) | 2101 (491) | 1602 (553) | 1902 (562) |
| RT (ms) | Pre | 294 (15) | 309 (32) | 305 (30) | 296 (17) | 302 (26) |
|  | Post | 297 (28) | 298 (32) | 301 (35) | 292 (19) | 298 (29) |

Duke = Duke University; IU = Indiana University Bloomington; DVT = Dynamic Vision Training; PVT = Placebo Vision Training; Pre = pre-training; Post = post-training; Vision Expectations = expected training benefit for vision; Performance Expectations = expected training benefit for performance; MOT = multiple-object tracking

**Table 2:** Means (and SDs if applicable) for measures of instrumented batting practice and NCAA game statistics

|  |  | Group |  | Team |  | All |
| --- | --- | --- | --- | --- | --- | --- |
|  |  | DVT | PVT | Duke | IU |  |
| Batting Practice |  |  |  |  |  |  |
| Number of Participants |  | 7 | 7 | 6 | 8 | 14 |
| Number of Training Session |  | 17.4 (8.3) | 17.1 (8.1) | 9 (2.4) | 23.5 (2.8) | 17.3 (7.9) |
| Perform Expectations | Pre | 4.43 (0.79) | 5 (0.82) | 5.33 (0.82) | 4.25 (0.46) | 4.71 (0.83) |
|  | Post | 4.86 (0.69) | 5 (1) | 5.5 (0.84) | 4.5 (0.53) | 4.93 (0.83) |
| Pitches Analyzed | Pre | 106 (77) | 126 (78) | 36.7 (19) | 176 (29) | 116 (75) |
|  | Post | 111 (86) | 113 (91) | 17.8 (4.2) | 182 (16) | 112 (85) |
| Exit Velocity (mph) | Pre | 87.6 (5.5) <sup>ψ</sup> | 87.7 (4.9) <sup>ψ</sup> | 91.9 (1.5) | 84.4 (4.2) | 87.6 (5) |
|  | Post | 88.7 (3.8) <sup>ψ</sup> | 88.7 (3.2) <sup>ψ</sup> | 89.3 (3) | 88.2 (3.7) | 88.7 (3.3) |
| Launch Angle (°) | Pre | 18.6 (7.8) <sup>ψ</sup> | 16.6 (5.9) <sup>ψ</sup> | 22.8 (7.4) | 13.7 (2) | 17.6 (6.7) |
|  | Post | 22.2 (6.3) <sup>ψ</sup> | 16.6 (7.5) <sup>ψ</sup> | 23.1 (9.3) | 16.7 (4) | 19.4 (7.3) |
| Batting Distance (ft) | Pre | 200 (47) <sup>ψ</sup> | 198 (53) <sup>ψ</sup> | 247 (26) | 163 (19) | 199 (48) |
|  | Post | 222 (28) <sup>ψ</sup> | 198 (28) <sup>ψ</sup> | 223 (30) | 201 (27) | 210 (30) |
| Season Statistics |  |  |  |  |  |  |
| Number of Participants |  | 5 | 7 | 6 | 6 | 12 |
| Number of Training Session |  | 19.8 (5.8) | 14.1 (8.5) | 13 (5.7) | 20 (8.4) | 16.5 (7.8) |
| Perform Expectations | Pre | 4.6 (0.89) | 4.71 (1.1) | 5.33 (0.82) | 4 (0.63) | 4.67 (0.98) |
|  | Post | 5.2 (0.45) | 5.17 (0.98) | 5.67 (0.52) | 4.6 (0.55) | 5.18 (0.75) |
| At-Bats | 2018 | 88.8 (86) | 90.4 (76) | 65 (81) | 114 (69) | 89.8 (77) |
|  | 2019 | 156 (86) | 164 (63) | 168 (81) | 152 (64) | 160 (70) |
| Batting Average | 2018 | 0.258 (0.062) | 0.284 (0.087) | 0.278 (0.099) | 0.269 (0.053) | 0.273 (0.075) |
|  | 2019 | 0.22 (0.084) | 0.258 (0.034) | 0.233 (0.077) | 0.252 (0.043) | 0.242 (0.06) |
| On-Base% | 2018 | 0.343 (0.07) | 0.374 (0.087) | 0.377 (0.11) | 0.344 (0.042) | 0.361 (0.078) |
|  | 2019 | 0.31 (0.1) | 0.355 (0.049) | 0.322 (0.09) | 0.351 (0.06) | 0.336 (0.075) |
| Slugging % | 2018 | 0.348 (0.14) | 0.397 (0.19) | 0.379 (0.22) | 0.375 (0.11) | 0.377 (0.17) |
|  | 2019 | 0.356 (0.17) | 0.379 (0.084) | 0.347 (0.14) | 0.392 (0.11) | 0.369 (0.12) |
| Walk % | 2018 | 0.0762 (0.061) | 0.0965 (0.047) | 0.0841 (0.071) | 0.092 (0.029) | 0.0881 (0.052) |
|  | 2019 | 0.102 (0.03) | 0.108 (0.031) | 0.0957 (0.025) | 0.115 (0.033) | 0.106 (0.03) |
| Strikeout % | 2018 | 0.326 (0.25) | 0.208 (0.058) | 0.322 (0.23) | 0.192 (0.045) | 0.257 (0.17) |
|  | 2019 | 0.308 (0.19) | 0.251 (0.045) | 0.307 (0.17) | 0.243 (0.04) | 0.275 (0.12) |

### 3.2. Testing Differential Expectation, Baseline Differences, and Training Session Adherence

Prior to and following training, participants were surveyed on their expectations regarding the potential benefits of the training program on both vision and performance. No group differences were identified on expectations of training benefits for vision at either the pre- (*p* > 0.75) or post- (*p* > 0.57) training periods, or in the change from pre- to post- training period (*p* > 0.34). Moreover, for the subsamples of subjects who completed batting practice (*n* = 14), no group differences were identified on expectations of training benefit for baseball performance at the pre- (*p* = 0.21) or post- (*p* = 0.76) training query, or in the change from pre- to post-training (*p* = 0.18). Similarly, for those participants with season statistics (*n* = 12), no group differences were identified on expectations of training benefit for baseball performance at either pre- (*p* = 0.85) or post- (*p* = 0.95) training, or in in the change from pre- to post- training (*p* = 0.17). This pattern of results is consistent with the intent-to-treat analysis. Overall, no evidence of differential expectations was found, and trainees generally indicated high expectation of training benefit on both vision and baseball performance throughout the training (i.e., ratings ≥ 4.7/6).

No group differences were identified on pre-training performance for any of the visual-motor variables (*ps* > 0.19), batting practice (*ps* > 0.87), or season statistics (*ps* > 0.25). This pattern of results is consistent with that from the intent-to-treat analysis. Thus, no evidence suggested a violation of group equivalence assumption on the tested variables at baseline. Training adherence was compared between the two training groups regarding the number of training sessions completed for each of the outcome-based subsamples. No significant difference of training adherence was identified between training groups in subsamples corresponding to visual-motor skill evaluations (*p* = 0.90), the instrumented batting practice (*p* = 0.95), or the NCAA season statistics (*p* = 0.23). This pattern of results is consistent with that from the intent-to-treat analysis. Overall no evidence suggested the groups had differential adherence to the training program.

### 3.3. Testing Generalized Near-Transfer

ANCOVA performed on the six visual-motor evaluation variables revealed no significant differences between training groups on Visual Clarity (*p* = 0.51), Contrast Sensitivity (*p* = 0.87), Near-Far Quickness (*p* = .71), Perception Span (*p* = 0.21), Multiple-Object Tracking (*p* = 0.39), or Reaction Time (*p* = 0.85). However, for all assessments except Visual Clarity, pre-training performance was a significant covariate (*ps* < 0.04) of the corresponding post-training performance in the ANCOVA model (see **Supplement 2.3** for the full ANCOVA output). These results were consistent with findings from the intent-to-treat analysis and suggest generalized visual-motor skills at the pre-training period had predictive power on their post-training assessments. They also suggest dynamic vision training did not transfer to improved generalized visual-motor skills relative to placebo vision training.

### 3.4. Testing Sports-Specific Intermediate Transfer to Instrumented Batting Practice Performance

The ANCOVA models fit reasonably well to the batting practice data combining individuals from the two training sites, with adjusted *R*^*2*^ values larger than 0.35 for Exit Velocity, 0.67 for Launch Angle, and 0.75 for Hit Distance. For each of the three batting practice variables, **Figure 4** descriptively shows the pre- and post-training group means (circles), standard deviations, and individual participant means (triangles), separately for the combined sample, the IU sample, and the Duke sample. Table 3 summarizes the ANCOVA results in detail.

**Figure 4.**
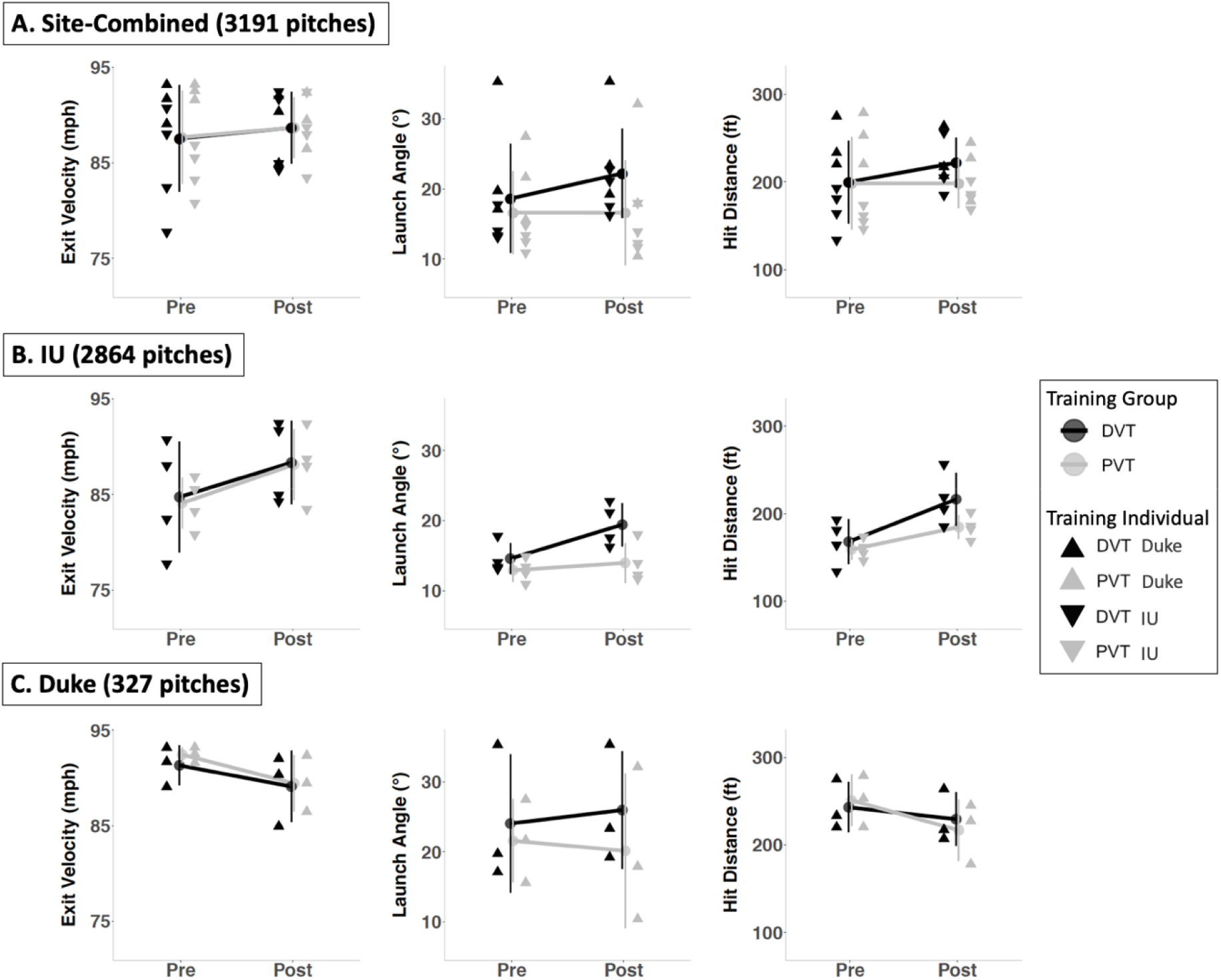
Descriptive illustrations of the group means (circles), standard deviations, and individual subject means (represented by triangles of two orientations and two colors differentiating individuals from the two teams and the training groups, respectively), at pre- and post-training periods for the three batting practice variables. Plots are shown with the number of total batting practice pitches observed for the (A) combined sample from both sites, (B) IU participants only, and (C) Duke participants only.

**Table 3:** ANCOVA model results for outcome measures of instrumented batting practice

| Combined | Exit Velocity |  |  | Launch Angle |  |  | Hit Distance |  |  |
| --- | --- | --- | --- | --- | --- | --- | --- | --- | --- |
|  | Estimate (SE) | t(9) | p | Estimate (SE) | t(9) | p | Estimate (SE) | t(9) | p |
| Training Group<sup>‡</sup> | 0.28 (1.65) | 0.17 | 0.43 | 9.38 (2.53) | 3.71 | 0.002** | 41.37 (9.10) | 4.55 | <0.001*** |
| Pre-Training Performance | 0.73 (0.23) | 3.17 | 0.01** | 0.63 (0.26) | 2.46 | 0.04* | 0.69 (0.19) | 3.54 | 0.006** |
| Performance Expectation | 0.37 (1.42) | 0.26 | 0.8 | 8.82 (2.28) | 3.87 | 0.004** | 33.21 (7.83) | 4.24 | 0.002** |
| Site | 4.92 (2.59) | 1.9 | 0.09 | 8.96 (4.52) | 1.98 | 0.08 | 70.84 (21.06) | 3.36 | 0.008** |
| IU | Exit Velocity |  |  | Launch Angle |  |  | Hit Distance |  |  |
|  | Estimate (SE) | t(4) | p | Estimate (SE) | t(4) | p | Estimate (SE) | t(4) | p |
| Training Group<sup>‡</sup> | 0.24 (2.36) | 0.1 | 0.46 | 5.32 (2.31) | 2.3 | 0.04* | 33.87 (15.01) | 2.26 | 0.04* |
| Pre-Training Performance | 0.75 (0.25) | 3.02 | 0.04* | 0.89 (0.51) | 1.75 | 0.16 | 0.86 (0.35) | 2.47 | 0.07 |
| Performance Expectation | 1.01 (2.76) | 0.37 | 0.73 | 2.64 (2.40) | 1.1 | 0.33 | 19.61 (17.03) | 1.15 | 0.31 |
| Duke | Exit Velocity |  |  | Launch Angle |  |  | Hit Distance |  |  |
|  | Estimate (SE) | t(2) | p | Estimate (SE) | t(2) | p | Estimate (SE) | t(2) | p |
| Training Group<sup>‡</sup> | 0.30 (3.43) | 0.09 | 0.47 | 11.71 (5.74) | 2.04 | 0.09 | 42.14 (14.27) | 2.95 | 0.05* |
| Pre-Training Performance | 3.76 (4.29) | 0.88 | 0.47 | 0.78 (0.44) | 1.8 | 0.21 | 0.58 (0.27) | 2.16 | 0.16 |
| Performance Expectation | -5.43 (8.29) | -0.65 | 0.58 | 11.73 (4.33) | 2.71 | 0.11 | 37.33 (9.55) | 3.91 | 0.06 |

Duke PVT, IU PVT, and Duke PVT serves as the reference group in the combined, IU, and Duke results, respectively. Training Group = DVT vs. PVT; Pre-Training Performance = the batting performance variable measured at pre-training; Performance Expectation = pre-training expectations of training benefit on batting performance; Site = Duke vs. IU. <sup>‡</sup> t-tests associated with the parameter are one-sided, whereas all others are two-sided. \* $p < .05$ , \*\* $p < .01$ , \*\*\* $p < .001$

Results from the combined site analysis revealed that, when controlling for pre-training performance, performance expectation, and site, the DVT group showed significantly greater improvement in Launch Angle (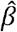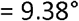, *t*[9] = 3.71, *p* = 0.002, Cohen’s *d* = 0.74) and Hit Distance (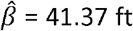, t[9] = 4.55, p < .001, Cohen’s *d* = 0.70), but not Exit Velocity (*p* = 0.87), relative to the PVT group. These inferential gains of 9.38° for Launch Angle and 41.37 ft for Hit Distance derived from the ANCOVA model correspond to descriptive gains of 3.6° for Launch Angle and 22 feet for Hit Distance when other factors are not considered (see **Figure 4A** and **Table 2**).

Pre-training performance was a significant covariate (*ps* < 0.04) of corresponding post-training performance for each of the three batting practice variables. In addition, performance expectations significantly predicted post-training performance on Launch Angle (*t*[9] = 3.87, *p* = 0.004) and Hit Distance (*t*[9] = 4.24, *p* = 0.002) and there was a significant effect of site for Hit Distance as IU batters produced longer distances than Duke batters (*t*[9] = 3.36, *p* = 0.008), when controlling for other factors. The effect of training group was replicated on both Launch Angle (*p* = 0.003) and Hit Distance (*p* < 0.001) in the intent-to-treat analysis results (**Supplement 2.1**).

In addition, because of differences in the equipment and procedures for the instrumented batting practice between the two sites, data from the two sites are analyzed separately. For the 2864 total batting practice observations among the 8 Indiana University (IU) athletes, ANCOVA results revealed that, when controlling for pre-training performance and performance expectation, the DVT group showed significant improvements in Launch Angle (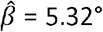, *t*[4] = 2.30, *p* = 0.042, Cohen’s *d* = 1.39) and Hit Distance (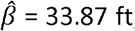, *t*[4] = 2.26, *p* = .043, Cohen’s *d* = 1.44), but not Exit Velocity (*p* = 0.46) relative to the PVT group. For the 327 total batting practice observations among the 6 Duke athletes, ANCOVA controlling for pre-training performance and performance expectation, demonstrated that the DVT group produced significantly longer Hit Distances (= 42.14 ft, *t*[2] = 2.95, *p* = .049, Cohen’s *d* = 0.54), relative to the PVT group, with non-significant differences for Launch Angles (= 11.71°, *t*[2] = 2.04, *p* = 0.089, Cohen’s *d* = 0.63) or Exit Velocity (*p* = 0.47).

### 3.5. Testing Sports-Specific Far Transfer to NCAA Game Statistics

An ANCOVA for the NCAA game statistics was run by weighting observations according to the number of at-bat in the 2018 season for the 12 athletes who had game statistics for the two seasons. These analyses res lted in non-significant differences between training groups on all five season statistics (see **Supplement 2.3** for the full ANCOVA output), including Batting Average (*p* = 0.91), On-Base % (*p* = 0.80), Slugging % (*p* = 0.69), Walk % (*p* = 0.29), and Strikeout % (*p* = 0.75). These results were consistent with those from the intent-to-treat analysis and did not change when the observations were otherwise weighted using the number of at-bats in the 2019 season, or the average of the 2018 and 2019 seasons.

## 4. Discussion

Vision is central to baseball, and there has long been a desire to improve batting performance through the application of vision-based training programs. However, there is scarce evidence demonstrating that such training may lead to performance benefits. This paper presents a first-of-kind, pre-registered, randomized, and placebo-controlled study attempting to determine whether the introduction of dynamic vision training drills, relative to placebo drills, leads to improvements in sports-specific performance and generalized visual-motor skills. Conceptually guided by the Modified Perceptual Training Framework and the Content-Context Transfer Taxonomy, the study leverages new opportunities arising with digital training platforms in a multi-week training protocol implemented with batters from two collegiate baseball teams. The training protocol generated a high adherence rate (83.33% completion rate with an average of 17 training sessions per person) with no difference between the two groups for expectations. While no training group effects were identified for generalized near-transfer (visual-motor skills) or sports-specific far-transfer (season game statistics), both the complete-case and the intent-to-treat analyses revealed training-induced enhancement in batting practice that signals sports-specific intermediate transfer. In particular, according to the pre-registration analysis plan, ANCOVA incorporating pre-training performance, expectations and site demonstrated significant group effects for Launch Angle and Hit Distance for DVT over PVT, amounting to medium-to-large effect sizes according to Cohen’s standards (*d* = 0.74 and 0.70, respectively). This corresponds to descriptive gains of 3.6° for Launch Angle and 22 ft for Hit Distance when other factors are not considered (see **Figure 4A** and **Table 2**).The intent-to-treat analysis also revealed consistent training effects, supporting the robustness of the findings against the existing attrition in the sample. While these findings are based on relatively small sample sizes ranging from 12 to 20 individuals, the presence of positive transfer and the absence of negative transfer support the potential utility of this type of vision training program in applied settings beyond baseball. Moreover, the placebo-controlled design used here offers a model for future studies aiming to test the transfer of skill training to high-performance contexts while also contributing to understanding of mechanisms underlying visual-motor learning.

In the following sections we discuss the training program and design considerations in this study, transfer effects that were (not) observed, limitations in this study, and future directions that might be most profitable for informing future studies.

### 4.1. Training Program and Study Design

Skilled performance in most athletic activities requires a combination of foundational visual skills, such as acuity and depth perception [14, 55], as well as general perceptual-cognitive skills such as anticipation, sustained attention, and decision-making [56]. Broadly speaking, training programs have been developed to improve these skills separately. Sports vision training approaches aim to enhance foundational visual skills utilizing generic visual stimuli, while perceptual-cognitive training aims to sharpen sports-specific visual skills through video or imagery presented in spatial or temporal contexts that mimic the sport. While the objectives and approaches are different, techniques across both of these domains are based on the key assumptions that (1) the targeted skill should differentiate athletes of different skill levels, (2) skill improvements should be possible through training, and (3) these improvements will transfer to enhance on-field performance [41].

While there is growing evidence supporting benefits of many visual and perceptual training techniques, the current study sought to integrate approaches across the spectrum of these dimensions into a comprehensive Dynamic Vision Training program that includes many of the most promising individual training drills. Building on past research with tablet-based vision training [16], anticipatory timing training [57, 58], and stroboscopic visual training in simplified [25, 26, 29] and complex environments [27, 28], the current study assembled these techniques into a brief, adaptive training circuit. The goal of this DVT circuit was to offer diverse, challenging, and engaging training activities possessing a high correspondence in the MPFT framework with the sports-defined skillset of interest — baseball batting.

To improve upon past studies that have tested sports-vision training in applied contexts [16, 36, 42], the current study employed a randomized and placebo-controlled design in order to balance exposure and training effort between the active and control participants. A key feature of this design was the implementation of placebo training activities in which participants performed challenging training drills that were modified so as not to engage the targeted skill elements, such as by training depth perception monocularly so as to remove the binocular depth cues and negate benefits of vergence training. Moreover, to balance effort and engagement, tasks were also conducted on adaptive schedules, leading to sustained engagement and equal compliance and adherence relative to the active training group.

### 4.2. Potential Positive Transfer to Batting Practice

A key objective of this study was to assess for transfer of training to sports-specific performance; therefore, the observation that DVT training may have led to improved batting practice measures supports one of the central hypotheses of this study, as articulated in the pre-registration. While past studies have provided evidence of transfer benefits [16,42], they generally have not utilized rigorous interventional designs such as the current one, thus leading to weaker evidence. Comparatively, the strength of this study includes the observed effects, consistency between the complete-case and intent-to-treat analyses, pre-registration of hypotheses and designs, a standardized vision training program between the two sites, and measuring batters’ training expectations [39].

Importantly, the possible transfer to batting performance is meaningful in an applied context. As indicated by the medium-to-large effect sizes observed in both launch angle and hit distance according Cohen’s standards [59], the DVT group demonstrated gains in these two important performance measures that can heavily impact outcomes in competitive games. In particular, there has been a recent emphasis on increasing batters’ launch angle for optimal batting performance, pointing to the importance of this skill. Overall this intervention approach is also effective when considering that, on average, each batter spent a total of 8.5 hours training with the protocol. A potential mechanism of transfer is that the training enhanced the skill elements critical for hitting a pitch with the bat’s ‘sweet spot’ [60]. This rationale can also explain the absence of transfer to exit velocity in batting performance because exit velocity may require skill elements relying more on power or strength, instead of vision. Future studies are encouraged to test such an explanation.

### 4.3. Null Training Results, Limitations and Future Directions

While the ultimate goal of this, and any other, training study is to transfer learning to game performance, a far transfer was not observed in game statistics. This null result can be explained by several possible factors. First, game performance can be viewed as a particularly distant transfer of training, which is harder to “simulate” along the MPTF dimensions. Although some training drills were purposefully designed to be sports-specific, it is challenging to simulate the physical, social, strategic, and functional context of actual collegiate baseball games. Second, even if some transfer exists from training to game statistics, such a “signal” may be hard to detect due to noise contributed by factors such as the defensive performance of opponents or coaching decisions. These sources of noise cannot be adequately controlled for using the current data, thereby masking potential training gains. Finally, the sample of athletes with pre- and post- training game statistics was relatively limited, consisting of 12 athletes who provided both pre- and post- training game statistics, with variable numbers of at bats for each season.

No generalized near transfer was observed on visual-motor skills measured on the Senaptec Sensory Station Tablet. The lack of such generalization may be explained by CCTT in that, although the transfer distance is considered “near” because batters practice with the digital tools in training, the amount of shared skill elements is small between the trained set of skill elements and the measured generalized visual-motor skills, suggesting visual-motor skill elements may be rather independent. This null finding also helps understand the lack of transfer findings from the generalized vision approaches in that training of general visual skills (“vision hardware”) is not likely to enhance specific ones that underpin the transfer of training to sports-specific performance. Nevertheless, both the training and placebo groups demonstrated pre-to-post improvements on all the generalized visual-motor skill measures, implying a learning effect. Such a learning factor demands attention from future researchers intending to gauge sports performance with digital tools.

The current study was designed to match rigorous interventional standards, while maximizing data collection from athletes who generally have limited time for research activities. The result is a well-controlled design, although it is still limited in several ways. Most notably, this study was conducted in the fall, between NCAA seasons, with all available and willing varsity baseball batters at two universities. This led to a sample of 20 individuals who completed an average of 8.5 hours of training in the generalized near transfer measures, 14 who completed the sports-specific intermediate transfer measures, and 12 with sufficient game statistics to evaluate sports-specific far transfer. While this sample is on par with previous training studies [16, 36, 42], there is still limited inferential power from a sample of this size. Future studies should aim to acquire multiple seasons and larger sample sizes. A second limitation resides in the instrumented batting practice data. Specifically, the Duke data were collected in the context of the baseball team’s typical training activities during the period of the study. Although these data can be regarded as more realistic measures, only a limited number of observations were available for athletes at this site. Thus, analyses based on separate data from Duke and IU, respectively, were conducted, in addition to the planned combined ANCOVA analyses. Nevertheless, the pattern of results were consistent with the inference that vision-based training transfers to batting practice performance, suggesting to some degree the robustness of the finding.

### 4.4. Conclusion

Following the combined frameworks of MPTF, TIE, and CCTT, the present training study with collegiate batters is the first to include causal evidence supporting training-induced, sports-specific transfer to baseball batting performance. The results invite ensuing training studies to improve upon the design and utilize new and emerging visual training tools to further investigate this emerging area of skill training and transfer.

## Declaration Page

### Ethics approval

The study was approved by both Duke University and Indiana University Bloomington research ethics committees, and the authors certify that the study was performed in accordance with the ethical standards in the 1964 Declaration of Helsinki and its later amendments.

### Consent to participate

All the participants in the study gave informed consent for both research participation and publication of their data.

### Consent for publication

All authors agreed with the content and that all gave explicit consent to submit.

### Availability of Data and Material

The research data and materials will be available upon requests.

### Code availability

The data processing code in R will be available upon requests.

### Authors’ contributions

Conceptualization: L. Gregory Appelbaum, Sicong Liu, Nicholas Port, Lyndsey M. Ferris; Methodology: Sicong Liu, L. Gregory Appelbaum, Lyndsey M. Ferris, Nicholas Port, Susan Hilbig, Edem Asamoa, John L. LaRue, Don Lyon, Katie Connolly; Formal analysis and investigation: Sicong Liu, Nicholas Port; Writing - original draft preparation: Sicong Liu, Lawrence Gregory Appelbaum, Susan Hilbig, Lyndsey M. Ferris, Edem Asamoa, John L. LaRue; Writing - review and editing: Sicong Liu, L. Gregory Appelbaum, Lyndsey M. Ferris, Nicholas Port, Susan Hilbig; Funding acquisition: L. Gregory Appelbaum.

## Acknowledgments

The authors would like to thank both the Duke and Indiana University baseball teams for participating in this research study. In particular, we would like to thank the players who showed great willingness to participate in our research and the coaches for facilitating their participation. The authors would also like to thank Christopher Gordon, Erikson Nichols, Kelsey Rankin, Fred Edmunds, and Kyle Burris for their valuable contributions to this study, as well as Paula Kelbly, Katie Heeter, Oliver Hobson, Jacob Thayer, Brett Miller, and the Indiana University School of Optometry professional students for their dedication to data collection and training. Lastly, the authors greatly appreciate the photography and video support of the Indiana University and Duke University Communications.

## Footnotes

a As IU assessed each batter five times both at pre- and post- training periods, IU assessment scores on a given visual-motor task at pre- and post- training periods was represented by averaging the the pre- and post- training assessed values, respectively.

