## Supplement 1 for "Dynamic Vision Training Transfers Positively to Batting Performance Among Collegiate Baseball Batters"

**Supplement 1: Methodological Details of Training Procedures**

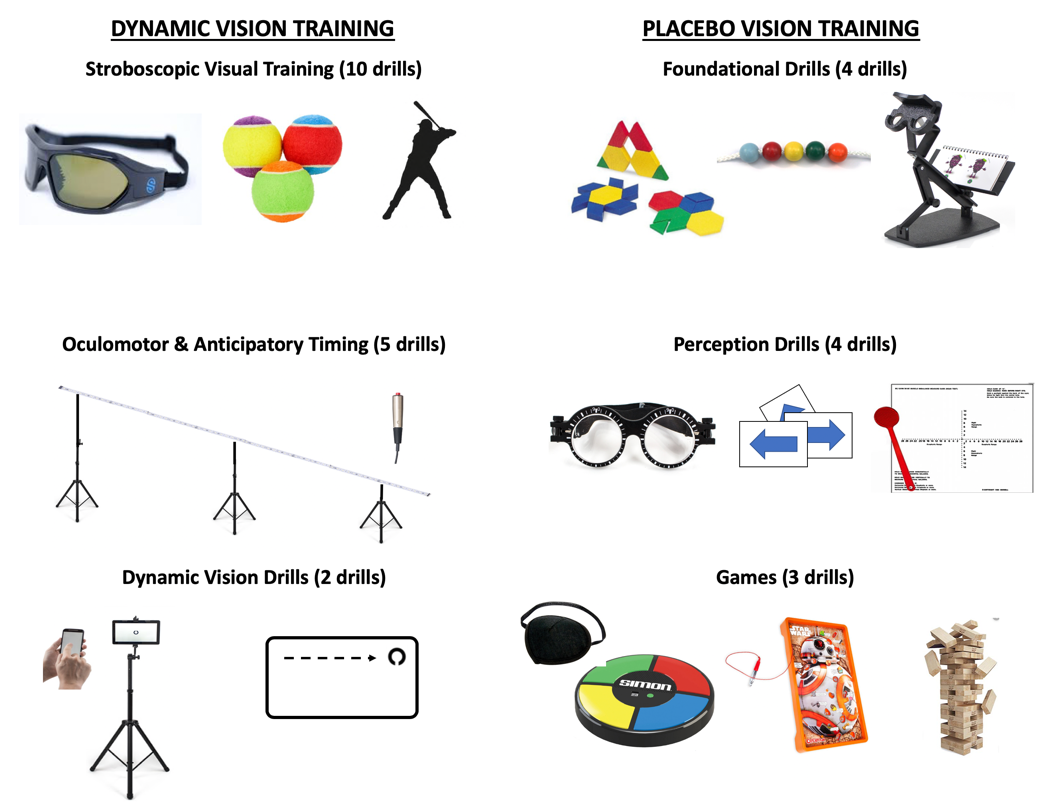

***Figure S1 (copied from manuscript)****. Illustrations and examples of the Dynamic Vision Training or Placebo Vision Training activities. DVT consisted of 10 different Stroboscopic Visual Training drills, which involved wearing strobe eyewear during catching and hitting drills; Oculomotor and Anticipatory Timing drills, which required participants to track or respond to fast moving lights on a light rail; and Dynamic Vision drills done on a digital tablet. PVT consisted of Foundational Drills including vertical oculomotor stability, pattern recognition, and attention to detail; Perception drills with adapted visual therapies; and Games, which incorporated games such as Simon, Operation, and Jenga with visual challenges such as eye patches or diopter lenses. PVT drills were modified such as not to directly engage or train horizontal oculomotor skills, visual-reaction timing, or depth/field of view perception.*

**1. Digital Vision Training**

DVT consisted of a regimen of up to 17 drills utilizing Senaptec training devices. Each session was divided into thirds, allowing for periods of time with the Sensory Station Tablet, Synchrony light rail, and Strobe glasses. Time spent with each device varied as participants progressed through training, but each session totaled 30 minutes.

In a typical session, a participant would start with the Senaptec strobe goggles for an assortment of 5 drills. Warm-up would consist of tossing a tennis ball to oneself and/or the experimenter with the goggles in an off position or easier setting, depending on the drills completed previously. The participant would then incrementally adjust the strobe difficulty within each drill, moving up after a quota of clean catches had been achieved. Similar means of progression were used for all sections and drills.

The second section involved use of the Synchrony light rail for 5 drills. Participants progressed through drills training reaction time and smooth/saccadic tracking. The third section used the Sensory Station Tablet for 2 drills testing near/far vision and contrast sensitivity. Advanced participants who achieved performance at or above level six in the initial 5 strobe drills had the opportunity to conclude each session by returning to the Senaptec strobe glasses and applying the glasses’ interference to a set of batting drills.

**1.1. Stroboscopic Drills – Ball Handling and Batting Cage (10 drills)**.

1.1.1. Ball Handling (8 drills) - The participant wore the Senaptec strobe glasses while doing an assortment of hand-eye coordination drills indoors. (1) The *warm-up* drill began by having the participant toss a tennis ball in the air and catch it with the strobe glasses set to “off” for approximately 30 seconds. (2) Starting at level one, the between-hand drill asked the participant to toss the ball from one hand to the other for approximately 30 seconds. (3, 4) In the separate-hand drill, the participant moved to tossing and catching with the right and left hand individually, each for approximately 30 seconds. (5, 6) In the between-person drill, the test administrator tossed the ball to the participant, targeting each hand individually until 5 number of catches had been achieved. (7, 8) Finally, in the turn drill, the participant would start with his back to the administrator and listen for a “ball” call signaling that he should jump and pivot 180 degrees to make the catch, finishing each trial by facing the administrator. This was deemed completed when 5 successful catches had been achieved on each side.
For all drills, difficulty varied by increasing the strobe glasses resistance and creating a longer period of blackout (for example, level three consists of 100 ms transparent to 150 ms opaque). Increases in difficulty occurred after five consecutive catches at the present level. As the participant progressed, they began each session closer to their highest strobe level and most recent set of drills.

1.1.2. Batting Cage (2 drills) - Participants who completed the initial strobe baseball progression by consistently reaching level six in all 8 previous drills advanced to training in the batting cages. (1) In the tee-ball batting drill, the participant began by tee-batting tennis balls with the strobe glasses set to off. Then, starting at level one, the participant batted with interference from the goggles and increased resistance as he progressed. If he achieved five consecutive hits at ideal exit angle (approximately 42 degrees), straightness, and velocity, he would increase the strobe glasses level. (2) In the tossed-ball batting drill, the administrator would stand behind a net shield and toss a baseball to the participant. Criteria for progression remained the same as the first drill.

**1.2. Senaptec Sensory Station Tablet (2 drills)**.

For both tablet drills, the tablet screen was positioned at eye level, 10ft from the participant. The participant used a handheld smartphone remote to log responses.

1.2.1. Dynamic Visual Acuity. In these drills three Landolt rings (semicricles with directional openings) would display on a tablet; the openings could all be in the same direction, or one ring could be out of the phase from the others. The participant was asked to swipe to the direction of the most-occurring opening (2/3 or 3/3 openings). The task difficulty adjusted through changes in the opacity, phase, and size of the rings.

1.2.2. Near-Far Quickness. In the near-far quickness drill, participants were instructed to hold the smartphone remote at approximately arm’s length and eye level, visually in line with the tablet. Landolt rings were alternately presented, one at a time, on either the tablet or the remote. Participants were asked to shift their focus as quickly as possible from one screen to the other, providing a response by swiping in the direction of the ring opening as in the previous drill. Task difficulty adjusted through changes in the size and rate of the rings presented.

**1.3. Synchrony Light Rail (5 drills)**.

The Synchrony light rail consists of a strip of 100 LEDs embedded in an 18-foot long flexible wire. The synchrony was supported by custom built PCV pipe frame that was designed to mimic the flight path of a pitched baseball. At the far end of the frame the height was 84 inches above the ground (approximately the release point of a pitched baseball) sloping down to the near end which was 40 inches above the ground (approximately the center of the strike zone). The far end represented the origin of all light sequences, while the near end represented the terminus and included the touch-sensitive response trigger.

For the first two drills, the participant stood perpendicular to the light rail and five feet back from its center, with the higher end on the right and the lower end on the left. Without moving his neck, the participant would track the light with only his eyes. (1) In the Smooth-Pursuit drill, the light would flow back and forth along the light rail. (2) In the Saccadic drill, a single light would illuminate at a time and jump to an unpredictable point in either direction along the light rail. In both drills, the difficulty varied based on speed of the light.

For the latter three drills, the participant moved to stand at the lower, near end of the light rail and held the touch-sensitive trigger in a comfortable batting stance while gazing along the light rail. All of these drills depended upon the response of the participant by pressing the trigger while the light flowed down the rail. (3)

In the Timing drill, a point 10 inches from the end of the light rail was indicated with a glowing sticker. The participant was asked to press the trigger when the light reached this point. (4) In the Go/No-Go drill, the light progressed in the same manner. However, participants were instructed to withhold response in the event that the light changed color from white to red at any point along the light rail. This color change would occur in only half the trials, and could occur at any point along the light’s path but was permanent once implemented. (5) In the Chase drill, a red light and a white light of variable speeds would race along the light rail toward the participant after starting at separate origins. The participant had to respond at the moment the two lights intersected. The difficulty for all three drills varied through increased and/or randomization of speed.

**2. Placebo Vision Training**

PVT consisted of a regimen of up to 11 drills utilizing vision therapy training methods that had been rendered ineffective due to the lack of therapeutic adjustment necessary. Each drill proceeded through phases of increasing difficulty. In general, the completion of four sessions of a drill led to the advancement to the next phase. Exceptions include the Thorington test and yoked prism (which were only included for 3 sessions), stick and straw drills (participants alternately covered one eye and were encouraged to exceed their previous score), and games (participants alternated whether Jenga or Operation was played).

**2.1. Foundational Drills (5 drills)**

The goal of these drills was to provide a consistent metric against which the participants could compete in order to capture their interest and provide rigor which might otherwise be lacking in contrast to the DVT. Therefore, these drills were included as components in nearly every visit and thus progressed through designated phases.

2.1.1. Stick and Straw Drill. In this drill the participant held a thin stick, covered one eye, and stood facing an administrator who held a straw. For a period of one minute, the administrator would present the straw from various positions and angles approximately 15 inches away. The participant attempted to insert the stick into the straw as quickly as possible, tallying as many alignments as possible in 60 seconds. As each phase advanced, the participant would alternate which eye was covered, and the administrator would progress from a static straw to a slowly, vertically-moving target.

2.1.2. Ductions Drill. Here, the participant wore an eyepatch over their non-dominant eye and was instructed to stand approximately 10 feet from administrators and fixate on a target as it was slowly moved. cover one eye at a time. The administrator stood holding a card displaying a black arrow on a white background, and would slowly move the card in a cross pattern, from the center out to the left, right, up, and down, passing back through the center each time. This pattern would be presented at approximately 15 inches from center, at arm’s length from center, and then again at arm’s length from center but standing at approximately 6 feet from the participant.

2.1.3. Bernell-O-Scope Drill. In this drill the participant sat a comfortable distance from the scope (Bernell, Inc, Mishawaka, IN; <https://www.bernell.com/product/BC200/Assessment_Kits>), and traced or drew Chieroscopic targets for five minutes, throughout which he received consistent reminders to draw as precisely as possible. Each phase utilized one eye at a time, with the corresponding hand being used to hold the pen. In phase one, the participant looked through the scope and traced the image oriented both correctly and upside down. In phase two, the participant replicated the figure freehand beside the present image. In phase three, the participant wore 2.0 diopter fogging lenses and traced the image. In phase four, the participant wore the same lenses and traced the image upside down.

2.1.4. Brock String. In the Brock string drill, the participant completed two-minute tasks of focusing on five different-colored beads tied along a string between 1 and 5 feet away from his eyes. One eye was patched at a time and the Brock string was held to the end of the player’s nose. In phase one, the participant was asked to alternate his focus from the close beads to the far beads every ten seconds. In phase two, participants were asked to follow the administrator’s pointer along the string, smoothly tracking along the length of the strength and alternating focus between the different beads. In phase three and four, participants continued these activities while using 2.0 diopter placebo yoked prism flippers.

2.1.5. Modified Thorington drill. Here, the participant stood approximately 5 feet from the administration, who stood holding a Thorington phoria card (Bernell, Inc) (<https://www.bernell.com/product/BC1209/Phoria-Tests>). The administrator directed a pin light through the test card while the participant held a Maddox rod up to one eye at a time, patching the other eye. The participant would begin with the Maddox rod oriented at 180^0^ and would describe the location her perceived the red line crossed the vertical axis of the Thorington card. The participant would then rotate the Maddox rod and describe the location he perceived the red line to cross the horizontal axis.

**2.2. Perception Drills (3 drills)**.

These drills utilized vision therapy tasks which challenge the eyes’ unity, or teamwork. They were implemented intermittently throughout the sessions.

2.2.1 Parquetry Blocks drills. Here, the participant sat at a desk with a parquetry pattern workbook (Bernell, Inc; <https://www.bernell.com/product/VTP/Games>) and blocks in front of him for five minutes each session. The participant assembled patterns of increasing complexity and decreasing guidance based upon the pattern book and progressed based upon his proficiency. During phase one, the participant was given the blocks and asked to reproduce patterns directly on top of the book’s pages. During phase two, the participant moved to completing patterns on a table surface next to the book. If successful, the participant continued to replicate the patterns on a table surface while looking through a 2.0 yoked prism diopter flipper. Following the completion of each pattern, the participant was instructed to turn the flipper to force his eyes to refocus. During phase three, the participant continued completing patterns with the use of a 3.0 yoked prism diopter flipper with turns between each pattern. During phase four, the participant advanced to wear 2.0 diopter fogging lenses.

2.2.2. Optical Illusions Drills. In these drills participants were presented with a laptop displaying different visual illusions (<https://michaelbach.de/ot/index.html>). The lilac chaser illusion was presented for 30 seconds. This task presents a fixation cross within a circle of purple dots placed like numbers on an analog clock, with an omitted dot rotating around the circle. Participants were instructed to indicate any time all the purple dots disappeared. The motion induced blindness illusion was presented for the remaining 30 seconds. This task presents a fixation point and three yellow dots overlaid upon a spinning blue grid. Participants were again instructed to indicate any time all the yellow dots disappeared.

2.2.3. Yoked Prism Drills. In these drills the participants viewed a set of vision tests presented 10 feet away. Participants would patch their dominant eye and then read the letters on the vision test through a 2.0 yoked prism diopter flipper as they progressed. Administrators reminded participants to flip the lens in between each letter, and to progress as quickly as possible.

**2.3. Games (3 drills)**.

These tasks applied placebo vision therapy techniques to classic games.

2.3.1 Card Drills. In these playing card drills, participants started by playing a 4 card x 4 card memory game with Red/Green Playing Cards (Bernell, Inc) (<https://www.bernell.com/product/KEY4102B/Games>) while wearing polarized glasses. The objective was to pair red/green playing cards based on suit. In the next level, participants played the card game “War” against the administrator while wearing the polarized glasses and narrating the game.

2.3.2. Visual Memory Drills. In the visual memory drills, participants were presented with a laptop with the website mindgames.com (<https://www.mindgames.com/Memory+Games>) which has digital versions of memory games. Participants were given the option of playing one of four visual memory games (Dinosaur Eggs, Memory Challenge, Casino Cards Memory, or Simon Says). Participants would then play the respective game, using visual memory cues to either denote where specific targets were, or to reproduce visual color patterns. Game complexities increased as participants completed each level. Polarized lenses were introduced in phase two as the participants mastered each level.

2.3.3. Board Game Drills. These drills alternated between playing Jenga^TM^ and Operation^TM^ (Hasbro, Inc, Pawtucket, RI). The participant’s dominant eye was initially patched in phase one. In phase two, a 2.0 diopter fogging lenses was introduced, increasing the difficulty of the task. If the participant continued to complete the tasks without irritation from the lenses, he progressed to phase three, using 5.0 diopter foggling lenses while accomplishing the game’s activities.
