## Supplement 2 for "Dynamic Vision Training Transfers Positively to Batting Performance Among Collegiate Baseball Batters"

***Supplement 2: Supplemental Results***

***Supplement 2.1:***

***Intent-to-Treat Analysis Results***

Note: The organization of the intent-to-treat results below mirror that of the Results section in the manuscript. The specific section titles followed ‘S’ to distinguish from those of the Results section in the manuscript. In addition, the intent-to-treat analysis for season statistics includes an ANCOVA without having the season statistics from 2018 as a covariate because of the high proportion (i.e., 9 out of 21) of batters missing competitions in that season.

*Testing Differential Expectation, Baseline Differences, and Training Session Adherence*

Before and after training, participants (*N* = 24) were queried on their expectations regarding the potential benefits of the training program on both vision and performance. No group differences were identified on expectations of training benefit for vision at either the pre- (*p* > 0.99) or post- (*p* > 0.49) training periods, or at the change from pre- to post- training (*p* > 0.35). Moreover, no group differences were identified on expectations of training benefit for baseball performance either at pre- (*p* > 0.09) or post- (*p* = 0.46) training periods, or at the change from pre- to post- training (*p* > 0.09). Similarly, for those participants with season statistics (*n* = 21), no group differences were identified on expectations of training benefit for baseball performance at either the pre- (*p* = 0.29) or post- (*p* = 0.80) training periods, or at the change from pre- to post- training (*p* = 0.16). These results are consistent with those in the complete-case analyses. Overall no evidence of differential expectations was found, and trainees generally indicated high expectation of training benefit throughout the training.

Group differences at baseline (i.e., pre-training) were tested on variables of visual-motor assessment and batting performance (including batting practice and season statistics). No group differences were identified on variables of visual-motor (*p*s > 0.17) and batting practice (*p*s > 0.74). No intent-to-treat analysis was performed for season statistics because nine batters did not compete in the 2018 season. This pattern of results is consistent with that from the complete-case analysis. Thus, no evidence suggested a violation of group equivalence assumption on tested variables at baseline.

Group training adherence differences were compared for the number of training session completed for each of the outcome subsamples. No significant difference was identified between the athletes who completed the visual-motor skills evaluations (*p* = 0.76), the structured batting practice (*p* = 0.76), or the NCAA season statistics (*p* = 0.90). This pattern of results is consistent with that from the complete-case analysis. Overall no evidence suggested the groups had differential adherence to the training program.

*Testing Generalize Near Transfer*

No training group effects were identified in the ANCOVA results of the six visual-motor evaluation variables including Visual Clarity (*p* = 0.37), Contrast Sensitivity (*p* = 0.79), Near-Far Quickness (*p* = 0.30), Perception Span (*p* = 0.21), Multiple-Object Tracking (*p* = 0.35), or Reaction Time (*p* = 0.83). However, all pre-training assessments, except for Visual Clarity (*p* = .054), were significant covariates (*p*s < 0.01) of the corresponding post-training assessments in the ANCOVA models. This pattern of results was consistent with findings from the complete-case analysis. These results suggest that generalized visual-motor skills assessed at the pre-training period had predictive power on their post-training assessments, but that training the targeted visual skill elements did not transfer to participant’s generalized visual-motor skills.

*Testing Sports-Specific Intermediate Transfer to Structured Batting Practice Performance*

For the batting practice variables, ANCOVA results showed good fit to the model (i.e., adjusted *R^2^*s > 0.48) and revealed that, when controlling for pre-training performance, DVT group performed better than PVT group regarding Launch Angle, $\hat{\beta}$ = 6.09°, *t*(19) = 3.11, *p* = 0.003, Cohen’s *d* = 1.11, and Hit Distance, $\hat{\beta}$ = 30.95 ft, *t*(19) = 3.66, *p* < 0.001, Cohen’s *d* = 1.01, but not Exit Speed (*p* = 0.38). All pre-training variables were significant covariates (*p*s < 0.007) of the corresponding post-training measures in the ANCOVA models.

*Testing Sports-Specific Far Transfer to NCAA Game Statistics*

For the season statistics, the analyses resulted in non-significant training group effect on all the five season statistics, including Batting Average (*p* = 0.45), On-Base % (*p* = 0.53), Slugging % (*p* = 0.12), Walk % (*p* = 0.40), and Strikeout % (*p* = 0.21).These null results were consistent with those from the intent-to-treat analysis and suggest no evidence of trained visual skill elements transferring to NCAA game performance.

***Supplement 2.2:***

***Descriptive Tables for Team-Group Factorial Combinations***

|  |  | **Team-Group Factorial Combination** | | | | **All** |
| --- | --- | --- | --- | --- | --- | --- |
|  |  | **Duke-DVT** | **Duke-PVT** | **IU-DVT** | **IU-MVT** |  |
| Number of Participants | | 6 | 6 | 4 | 4 | 20 |
| Number of Training Session | | 12.7 (5.8) | 12.7 (4.9) | 24 (1.4) | 23 (4) | 17 (6.9) |
| ***Expectations*** | |  |  |  |  |  |
| Vision Expectations | Pre | 5 (0.63) | 5.67 (0.52) | 5 (0) | 4.25 (0.5) | 5.05 (0.69) |
|  | Post | 5.5 (0.84) | 5.83 (0.41) | 5.75 (0.5) | 4.75 (0.96) | 5.5 (0.76) |
| Perform Expectations | Pre | 5 (0.89) | 5.67 (0.52) | 4 (0) | 4.5 (0.58) | 4.9 (0.85) |
|  | Post | 5.17 (0.75) | 5.83 (0.41) | 4.75 (0.5) | 4.25 (0.5) | 5.1 (0.79) |
| ***Visual Motor Evaluations*** | |  |  |  |  |  |
| VC (LogMAR) | Pre | -0.0997 (0.11) | -0.166 (0.13) | -0.176 (0.038) | -0.199 (0.05) | -0.155 (0.1) |
|  | Post | -0.145 (0.14) | -0.153 (0.064) | -0.211 (0.063) | -0.226 (0.085) | -0.177 (0.096) |
| CS (log) | Pre | 1.57 (0.23) | 1.6 (0.22) | 1.55 (0.14) | 1.78 (0.3) | 1.62 (0.23) |
|  | Post | 1.6 (0.28) | 1.77 (0.34) | 1.53 (0.17) | 1.73 (0.14) | 1.66 (0.26) |
| NFQ (score) | Pre | 31 (4.7) | 27.5 (8.5) | 33.5 (7.5) | 32.1 (4.5) | 30.7 (6.5) |
|  | Post | 35.7 (8.8) | 33.3 (4.3) | 39.5 (2.6) | 36.6 (4.2) | 35.9 (5.8) |
| PS (score) | Pre | 40.5 (12) | 57.2 (8.5) | 51.6 (6.5) | 40.2 (13) | 47.7 (12) |
|  | Post | 46.3 (14) | 56.8 (6.6) | 56 (3.7) | 44.1 (9.9) | 51 (11) |
| MOT (score) | Pre | 1818 (355) | 2032 (368) | 1713 (174) | 1581 (753) | 1814 (438) |
|  | Post | 2141 (419) | 2062 (592) | 1579 (429) | 1625 (728) | 1902 (562) |
| RT (ms) | Pre | 295 (18) | 316 (38) | 294 (14) | 299 (22) | 302 (26) |
|  | Post | 297 (33) | 306 (39) | 298 (23) | 286 (15) | 298 (29) |

***Table S2.2.1:*** *Means (and SDs if applicable) for measures of participation, expectations and performance on the visual-motor evaluations. Duke = Duke University; IU = Indiana University Bloomington; DVT = dynamic vision training; PVT = placebo vision training; Pre = pre-training; Post = post-training; Vision Expectations = expected training benefit for vision; Perform Expectations = expected training benefit for performance; VC = visual clarity; CS = contrast sensitivity; NFQ = near-far quickness; PS = perception span; MOT = multiple-object tracking; RT = reaction time.*

|  |  | **Team-Group Factorial Combination** | | | | **All** |
| --- | --- | --- | --- | --- | --- | --- |
|  |  | **Duke-DVT** | **Duke-PVT** | **IU-DVT** | **IU-MVT** |  |
| ***Batting Practice*** |  |  |  |  |  |  |
| Number of Participants | | 3 | 3 | 4 | 4 | 14 |
| Number of Training Session | | 8.67 (2.1) | 9.33 (3.2) | 24 (1.4) | 23 (4) | 17.3 (7.9) |
| Perform Expectations | Pre | 5 (1) | 5.67 (0.58) | 4 (0) | 4.5 (0.58) | 4.71 (0.83) |
|  | Post | 5 (1) | 6 (0) | 4.75 (0.5) | 4.25 (0.5) | 4.93 (0.83) |
| At-Bats | Pre | 26.7 (7.1) | 46.7 (24) | 165 (29) | 186 (28) | 116 (75) |
|  | Post | 19.3 (3.8) | 16.3 (4.7) | 180 (12) | 185 (22) | 112 (85) |
| Exit Speed (mph) | Pre | 91.3 (2.1) | 92.5 (0.81) | 84.7 (5.8) | 84.1 (2.7) | 87.6 (5) |
|  | Post | 89.1 (3.7) | 89.4 (2.9) | 88.3 (4.3) | 88.1 (3.7) | 88.7 (3.3) |
| Launch Angle (°) | Pre | 24.1 (9.8) | 21.6 (5.9) | 14.6 (2.2) | 12.9 (1.6) | 17.6 (6.7) |
|  | Post | 26 (8.4) | 20.2 (11) | 19.4 (3) | 14 (2.9) | 19.4 (7.3) |
| Batting Distance (ft) | Pre | 243 (29) | 251 (29) | 168 (26) | 159 (12) | 199 (48) |
|  | Post | 229 (30) | 217 (35) | 216 (30) | 185 (14) | 210 (30) |
| ***Season Statistics*** |  |  |  |  |  |  |
| Number of Participants | | 3 | 3 | 2 | 4 | 12 |
| Number of Training Session | | 16.7 (5.5) | 16.7 (5.5) | 24.5 (0.71) | 17.8 (9.9) | 16.5 (7.8) |
| Perform Expectations | Pre | 5 (1) | 5 (1) | 4 (0) | 4 (0.82) | 4.67 (0.98) |
|  | Post | 5.33 (0.58) | 5.33 (0.58) | 5 (0) | 4.33 (0.58) | 5.18 (0.75) |
| At-Bats | 2018 | 39 (46) | 39 (46) | 164 (83) | 90 (58) | 89.8 (77) |
|  | 2019 | 138 (109) | 138 (109) | 182 (62) | 138 (68) | 160 (70) |
| Batting Average | 2018 | 0.228 (0.04) | 0.228 (0.04) | 0.302 (0.076) | 0.252 (0.039) | 0.273 (0.075) |
|  | 2019 | 0.207 (0.11) | 0.207 (0.11) | 0.24 (0.043) | 0.257 (0.048) | 0.242 (0.06) |
| On-Base% | 2018 | 0.341 (0.08) | 0.341 (0.08) | 0.346 (0.083) | 0.344 (0.026) | 0.361 (0.078) |
|  | 2019 | 0.297 (0.13) | 0.297 (0.13) | 0.331 (0.061) | 0.361 (0.066) | 0.336 (0.075) |
| Slugging % | 2018 | 0.272 (0.1) | 0.272 (0.1) | 0.463 (0.13) | 0.331 (0.093) | 0.377 (0.17) |
|  | 2019 | 0.326 (0.22) | 0.326 (0.22) | 0.401 (0.14) | 0.388 (0.11) | 0.369 (0.12) |
| Walk % | 2018 | 0.0851 (0.083) | 0.0851 (0.083) | 0.0628 (0.015) | 0.107 (0.021) | 0.0881 (0.052) |
|  | 2019 | 0.0949 (0.029) | 0.0949 (0.029) | 0.113 (0.04) | 0.117 (0.036) | 0.106 (0.03) |
| Strikeout % | 2018 | 0.44 (0.28) | 0.44 (0.28) | 0.154 (0.0039) | 0.211 (0.044) | 0.257 (0.17) |
|  | 2019 | 0.339 (0.26) | 0.339 (0.26) | 0.262 (0.036) | 0.233 (0.044) | 0.275 (0.12) |

***Table S2.2.2:*** *Means (and SDs if applicable) for measures of batting practice and NCAA game statistics. Duke = Duke University; IU = Indiana University Bloomington; DVT = dynamic vision training; PVT = placebo vision training; Pre = pre-training; Post = post-training; Perform Expectations = expected training benefit for performance.*

***Supplement 2.3:***

***ANCOVA Tables for Generalized Near Transfer and Sports-Specific Far Transfer***

|  | **Visual Clarity** | | |  | **Contrast Sensitivity** | | |  | **Near-Far Quickness** | | |
| --- | --- | --- | --- | --- | --- | --- | --- | --- | --- | --- | --- |
|  | Estimate (SE) | *t*(15) | *p* |  | Estimate (SE) | *t*(15) | *p* |  | Estimate (SE) | *t*(15) | *p* |
| Training Group*^ψ^* | 0.0009 (0.04) | 0.02 | 0.51 |  | 0.12 (0.10) | 1.19 | 0.87 |  | 1.38 (2.41) | 0.57 | 0.71 |
| Pre-Training Performance | 0.27 (0.23) | 1.19 | 0.25 |  | 0.60 (0.24) | 2.55 | 0.02^*^ |  | 0.45 (0.20) | 2.27 | 0.04^*^ |
| Vision Expectation | 0.04 (0.04) | 1.07 | 0.30 |  | -0.09 (0.09) | -1.05 | 0.31 |  | -0.09 (2.06) | -0.05 | 0.96 |
| Site | -0.03 (0.05) | -0.54 | 0.60 |  | -0.17 (0.12) | -1.45 | 0.17 |  | 1.88 (2.88) | 0.65 | 0.52 |

|  | **Perception Span** | | |  | **Multiple Object Tracking** | | |  | **Reaction Time** | | |
| --- | --- | --- | --- | --- | --- | --- | --- | --- | --- | --- | --- |
|  | Estimate (SE) | *t*(15) | *p* |  | Estimate (SE) | *t*(15) | *p* |  | Estimate (SE) | *t*(15) | *p* |
| Training Group*^ψ^* | 2.39 (2.86) | 0.84 | 0.21 |  | 63.49 (216.18) | 0.29 | 0.39 |  | 12.29 (11.64) | 1.06 | 0.85 |
| Pre-Training Performance | 0.77 (0.12) | 6.21 | <.001^***^ |  | 0.63 (0.27) | 2.35 | 0.03^*^ |  | 0.78 (0.24) | 3.32 | 0.005^**^ |
| Vision Expectation | -2.44 (2.50) | -0.98 | 0.34 |  | -135.09 (188.83) | -0.72 | 0.49 |  | 6.05 (9.70) | 0.62 | 0.54 |
| Site | -1.05 (3.32) | -0.32 | 0.76 |  | -420.43 (264.29) | -1.59 | 0.13 |  | 1.83 (13.44) | 0.14 | 0.89 |

***Table S2.3.1:*** *ANCOVA Table for Generalized Near Transfer.* *Duke PVT serves as the reference group. Training Group = DVT vs. PVT; Pre-Training Performance = batting performance variable measured prior to training; Vision Expectation = pre-training expectations of training benefit on vision; Site = Duke vs. IU. ^ψ^ t-tests associated with the parameter are one-sided, whereas all others are two-sided. ^*^ p < .05, ^**^ p < .01, ^***^ p < .001*

|  | **Batting Average** | | |  | **On-Base %** | | |  | **Slugging %** | | |
| --- | --- | --- | --- | --- | --- | --- | --- | --- | --- | --- | --- |
|  | Estimate (SE) | *t*(15) | *p* |  | Estimate (SE) | *t*(15) | *p* |  | Estimate (SE) | *t*(15) | *p* |
| Training Group*^ψ^* | -0.04 (0.03) | -1.46 | 0.91 |  | -0.04 (0.04) | 0.88 | 0.80 |  | -0.04 (0.07) | -0.53 | 0.69 |
| Pre-Training Performance | 0.43 (0.18) | 2.45 | 0.04^*^ |  | 0.49 (0.33) | 1.49 | 0.18 |  | 0.51 (0.22) | 2.36 | 0.05^*^ |
| Performance Expectation | -0.03 (0.02) | -1.47 | 0.19 |  | -0.03 (0.03) | -1.03 | 0.34 |  | -0.07 (0.05) | -1.41 | 0.20 |
| Site | -0.02 (0.03) | -0.77 | 0.47 |  | -0.009 (0.05) | -0.19 | 0.85 |  | -0.05 (0.07) | -0.62 | 0.56 |

|  | **Walk %** | | |  | **Strikeout %** | | |
| --- | --- | --- | --- | --- | --- | --- | --- |
|  | Estimate (SE) | *t*(15) | *p* |  | Estimate (SE) | *t*(15) | *p* |
| Training Group*^ψ^* | 0.01 (0.02) | 0.58 | 0.29 |  | 0.04 (0.06) | 0.72 | 0.75 |
| Pre-Training Performance | 0.30 (0.34) | 0.87 | 0.42 |  | 0.33 (0.45) | 0.72 | 0.49 |
| Performance Expectation | -0.002 (0.02) | -0.11 | 0.92 |  | 0.03 (0.04) | 0.59 | 0.58 |
| Site | 0.008 (0.02) | 0.32 | 0.76 |  | 0.0005 (0.07) | 0.01 | 1.00 |

***Table S2.3.2****: ANCOVA Table for Sports-Specific Far Transfer. Duke PVT serves as the reference group. Training Group = DVT vs. PVT; Pre-Training Performance = batting performance variable measured prior to training; Performance Expectation = pre-training expectations of training benefit on batting performance; Site = Duke vs. IU. ^ψ^ t-tests associated with the parameter are one-sided, whereas all others are two-sided. ^*^ p < .05, ^**^ p < .01, ^***^ p < .001*
